# The novel leishmanial Copper P-type ATPase plays a vital role in intracellular parasite survival

**DOI:** 10.1101/2021.01.01.425060

**Authors:** Rupam Paul, Sourav Banerjee, Samarpita Sen, Pratiksha Dubey, Saptarshi Maji, Anand K Bachhawat, Rupak Datta, Arnab Gupta

**Affiliations:** Department of Biological Sciences, Indian Institute of Science Education and Research Kolkata, Mohanpur, West Bengal -741246, India; Department of Biological Sciences, Indian Institute of Science Education and Research Mohali, Knowledge city, Sector 81, Manauli, PO, Sahibzada Ajit Singh Nagar, Punjab-140306, India

**Keywords:** ATP7, *Leishmania*, Copper, Cu-ATPase, lysosome, ATP7A, host-pathogen interaction

## Abstract

Copper is essential for all life forms; however in excess it is extremely toxic. Toxic properties of copper are utilized by hosts against various pathogenic invasions. *Leishmania*, in its both free-living and intracellular forms, was found to exhibit appreciable tolerance towards copper-stress. To determine the mechanism of copper-stress evasion employed by *Leishmania* we identified and characterized the hitherto unknown Copper-ATPase in *Leishmania major* and determined its role in parasite’s survival in host macrophage cells. *L. major* Cu-ATPase, LmATP7, exhibits high homology with its orthologues at multiple conserved motifs. In promastigotes, LmATP7 localized to the plasma membrane with a fraction in intracellular puncta. Upon copper treatment, *LmATP7* expression increases few folds. LmATP7 is capable of complementing copper transport in Cu-ATPase-Δ yeast strain. Promastigotes overexpressing *LmATP7* exhibits higher survival upon copper stress indicating efficacious copper export compared to wild type and heterozygous *LmATP7* knock-out parasites. We explored macrophage-*Leishmanial* interaction with respect to copper stress subjected by the host upon parasite and the parasite’s reciprocating response thereon to evade the stress. The Copper-P-type-ATPases ATP7A/7B serve as major copper exporters in mammals that maintain cellular copper levels. We found that *Leishmania* infection triggers ATP7A upregulation in macrophages. Additionally, as part of host response, ATP7A traffics from trans-Golgi network and transports copper to the endosomal and endolysosomal compartments harbouring the *Leishmania* amastigotes. Finally, we show *LmATP7* overexpression in this parasite increased amastigote survivability within infected macrophages whereas knocking it down reduces that drastically, establishing its role in combating host-induced copper stress.

## Introduction

*Leishmania* is a digenetic protozoan belonging to the trypanosomatid group that alternates between sandfly vector and mammalian hosts. They are known to cause a wide spectrum of tropical human diseases collectively known as leishmaniasis. Severity of the disease depends on the *Leishmania* species and is manifested by a range of symptoms that varies from disfiguring skin lesions to life-threatening infection of the visceral organs [1]. With more than a million new cases and 20,000-30,000 deaths every year, 12 million people are currently affected from about 100 endemic countries imposing a significant threat to the global healthcare system [2,3]. Once the flagellated promastigote form of *Leishmania* enter mammalian host via sand fly bite, they are rapidly phagocytosed by macrophages [4]. Phagocytosis occurs either directly or by engulfment of the parasite-harbouring apoptotic neutrophils followed by transformation from promastigotes into non-flagellated amastigotes in the phagolysosome. Within the acidic phagolysosomes, they continue to proliferate until the cell bursts, leading to the spread of the infection [5]. Inside this compartment, *Leishmania* has to withstand a variety of host-induced stress factors including free radicals, lysosomal hydrolases and low pH [6,7]. They are not only equipped to defend against such harsh environment but they can also manipulate host gene expression to their benefit [8,9,10]. Unavailability of a vaccine along with increasing resistance of the existing drugs makes it even more important to understand *Leishmania* physiology and the molecular mechanisms which allow them to thrive inside the host [11,12].

Copper is an essential micronutrient for biological system. Several enzymes, involved in catalysing biochemical processes, utilize the ability of copper to cycle between cuprous and cupric states. Shuttling between Cu(II) and Cu(I) states can lead to oxidative damage of cells via Fenton like reaction when free copper is available [13]. Organisms have evolved mechanisms where several proteins are involved in tightly regulating the bioavailability of copper [14]. Copper homeostasis, which includes regulating its transport and intracellular distribution, is crucial as excess copper can be detrimental. Copper binding proteins, chaperones, transporters keep intracellular free copper at a very low level in the order of 10^−18^M [15].

Various studies have shown that copper plays a key role in host-pathogen interaction. Copper-deficient hosts are more susceptible to several pathogens that include prokaryotes like *Salmonella typhimurium*, *Pasteurella hemolytica* and eukaryotes like *Candida albicans*, *Trypanosoma lewisi* [16,17,18,19]. The bactericidal activity of macrophages and neutrophils is also impaired upon copper deficiency [20,21]. Similarly, copper channelization to the phagosomal compartment of macrophage during *Mycobacterium avium* and *E.coli* infection indicated how hosts tend to utilize copper to fight off intracellular pathogens [22,23]. *E.coli* mutant with a defective copper exporting system showed significantly more susceptibility to copper mediated killing within macrophage [23].

The P-type Cu-ATPases are involved in removing excess copper from the cell and are one of the key players in maintaining copper homeostasis. In bacteria, CopA and CopB are the Cu-ATPases that carry out this function [24,25]. There is a single Cu-ATPase in lower eukaryotes (non-chordates) referred to as Ccc2p in yeast *Saccharomyeces cerevisiae* or ATP7 in *Drosophila melanogaster* or simply Cu-ATPase [26]. With increased complexity, ATP7A and ATP7B are branched out of ATP7 in higher eukaryotes [27]. In the present study, we have functionally characterized a novel copper transporting ATPase (LmATP7) of *L. major* (*LmATP7*) and determined its role in leishmanial survivability in host macrophage. Our study shows for the first time the physiological function of a full-length Cu-ATPase belonging to the kinetoplastida order. Additionally, our study reveals the protective role of LmATP7 in free living promastigotes in copper stress. Further, we also establish that intracellular amastigotes combat host-induced copper stress in lysosomes using LmATP7 that plays a key role in determining its pathogenicity and successful manifestation of infection.

## Results

### *Leishmania* genome encodes and expresses P-type Copper ATPase

Copper has been conventionally used as a chemotherapeutic agent against parasitic infections [22,23,28]. To explore if lethality could be induced in *Leishmania* parasites, we tested survivability of the parasite in growth media supplemented with copper (Cu) in a range between 0-500µM at 24, 48 and 72 hrs post incubation. At 50 and 100µM of external Cu concentration though the parasite showed a decline in its growth starting from 24 hrs, the trend was more drastic at 72 hrs post incubation (Fig. 1A). Importantly, at 200 and 300µM of Cu concentration there was no increase in parasite count at 24-72 hrs post incubation as compared to the initial inoculum. However, at 500µM of external Cu supplementation the parasite number gradually decreased with time starting from 24hrs post incubation indicating the toxic effect of Cu on the parasite. Interestingly, even at this high Cu concentration the number of promastigotes remained almost same at 48hrs and 72hrs post incubation (Fig. 1A). We were interested to further evaluate the effect of external copper on *Leishmania* survivability within macrophage cells. For this, we first infected J774A.1 macrophages with *L. major* promastigotes and further supplemented the growth medium with 0- 500 µM of Cu. Although we checked whether Cu has any toxicity over macrophage growth, we could not find any alteration in it when J774A.1 cells were treated with 0- 500 µM of external Cu for 0- 48hrs (Fig. S1). However, similar to our observation on copper induced toxicity over free living promastigotes we found the similar trend in intracellular amastigotes survivability with increasing copper concentration. As evident from Fig. 1B starting from 50 µM of Cu the parasite burden decreased significantly with about 3 folds reduction at 500µM external Cu as compared to untreated control post 24 hrs infection. However, given that the internal copper in the parasite is maintained at an extremely low level (∼2-4 ppb as measured in promastigotes using ICP-OES by us), the survivability trend indicated that *L. major* exhibits copper resistance or tolerance to an appreciable extent. We argue that an efficient copper handling machinery is present in the parasite that confers its survivability in elevated non-physiological copper.

**Fig 1.**
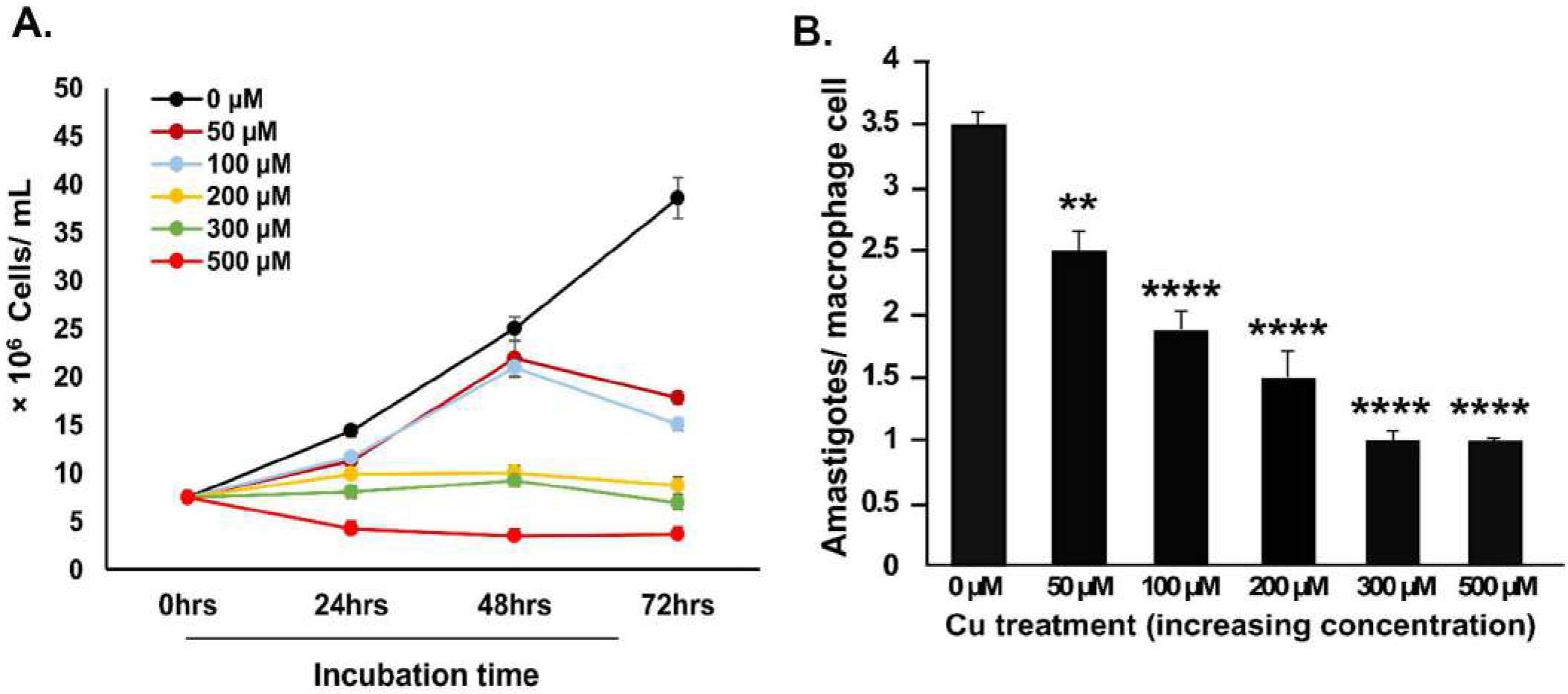
Survivability and growth of wild-type *L. major* promastigotes and amastigotes at increasing copper concentration. **(A)** Wild type *L. major* promastigotes were grown at 0- 500µM externally added Cu concentration for 24 hrs, 48 hrs and 72 hrs. Initially, 7.5×10^6^ cells were added to the media at time point 0 hrs and further at the above mentioned time-points cells were counted by haemocytometer-based cell counting. Error bars represent mean ± SD of values from three independent experiments. **(B)** J774A.1 macrophages were infected with wild type *L. major*. Bars represent amastigote count inside the macrophage at 0- 500µM of Cu concentration at 24 hrs post infection. Error bars represent mean ± SD of values from three independent experiments. * represents the significance of difference in number of amastigotes surviving inside macrophages under any external copper supplementation with respect to those surviving inside macrophages in absence of external copper. **P≤0.01, ****P≤0.0001 (Student’s t-test).

We attempted to identify if *Leishmania* genome encodes for any copper exporter that might be key for its survival in copper stress within the macrophage lysosomes during infection or in artificial experimental condition as tested in the previous section (as shown in Fig. 1A,B). We identified a putative copper transporting P-type ATPase (*LmATP7*) in the genome of *L. major* from the kinetoplastids informatics resources, TriTrypDB. It is predicted to have three heavy metal-binding domains (Pfam PF00430) in its N-terminal region, each containing the characteristic Cu^+^ binding motif, CXXC. It has the highly conserved TGE sequence in the actuator (A) domain and DKTGT sequence in the phosphorylation (P) domain. Both A and P regions are contained within the E1-E2 ATPase domain (Pfam PF00122), where E1 and E2 are the two cation binding states (Fig. 2A). The catalytic activity of Cu-ATPases is exhibited through phosphorylation and dephosphorylation of Asp residue in the conserved DKTGT motif. LmATP7 is also comprised of a hydrolase region (Pfam PF00702) containing TGDN and GXGXND conserved motifs in the ATP binding site of the nucleotide binding domain and the SX<u>H</u>P motif whose mutation in human orthologue ATP7B (His1069Gln) is the most frequent contributor to Wilson Disease in the Caucasian population (Fig. 2B). The SMART server predicted nine transmembrane domains (TMDs) in LmATP7, which is similar to the predicted TMDs in *Trypanosoma congolense* [29]. To date, no more than eight TMDs have been shown experimentally in Cu-ATPases across species. The extra TMD could be unique to some species of the kinetoplastida. LmATP7 contains copper coordinating CPC motif in the seventh TMD that is conserved in the IB P-type ATPase family. The genomic DNA and RNA of *L. major* were isolated and amplified (cDNA from RNA) using primers designed to amplify the *LmATP7*. Gel electrophoresis data showed a fragment of about 3500bp for both genomic DNA and cDNA indicating its transcriptional expression (Fig. 2C) which was subsequently confirmed by Sanger sequencing. The sequencing result revealed a DNA fragment of 3492bp, which is identical to the length of the putative copper-transporting ATPase-like protein encoding gene [*Leishmania major* strain Friedlin, GeneID 5654633]. Although both encoded identical amino acid sequence (1163 in length), *LmATP7* gene from our *L. major* strain 5ASKH differed in six nucleotides from the sequence of Friedlin strain. Both gene and mRNA sequence of *LmATP7* are submitted to the GenBank (accession numbers MW261996 and MW261995 respectively).

**Fig 2.**
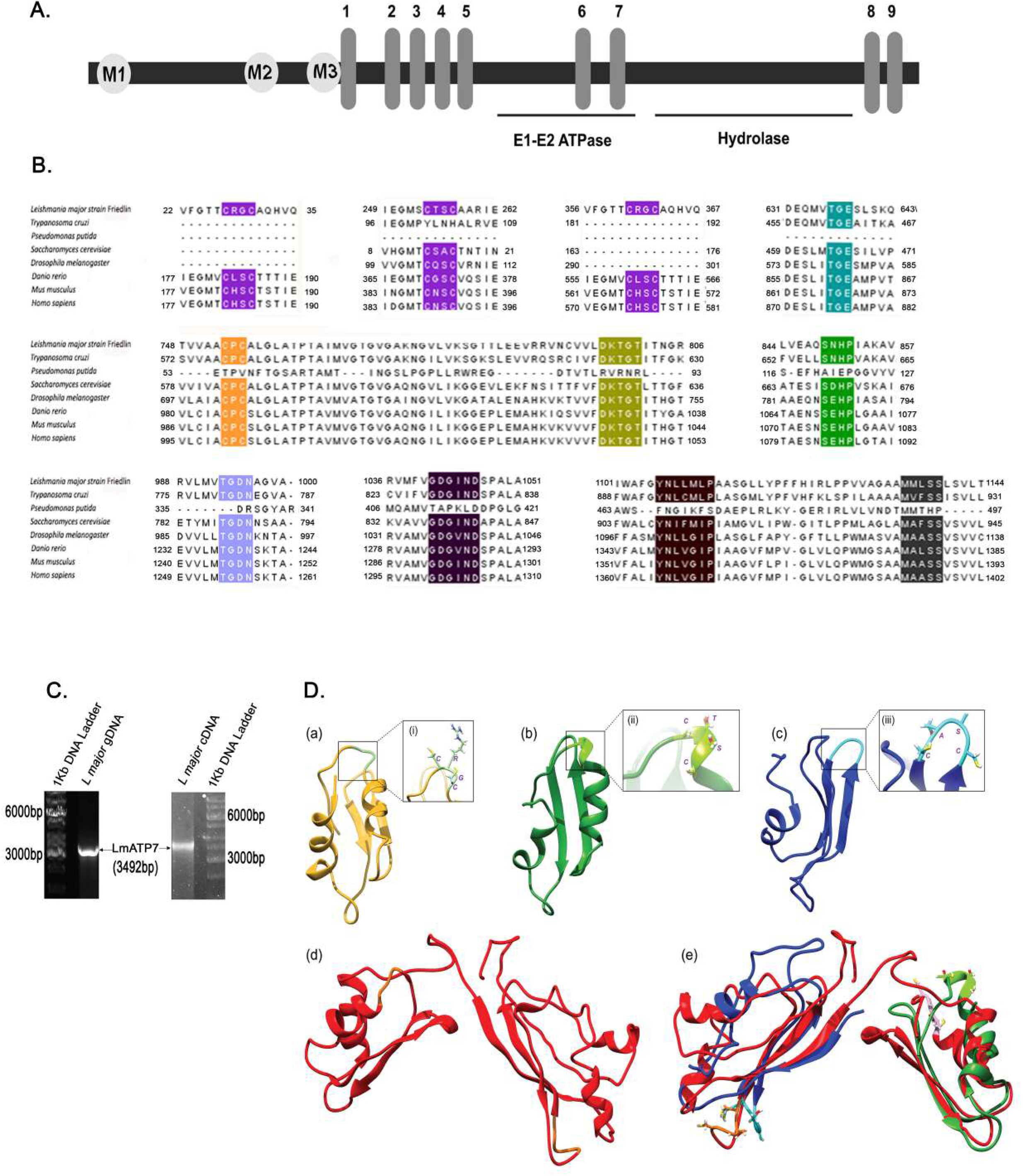
LmATP7 is a putative P-type copper ATPase. **(A)** Schematic representation of the putative Leishmanial copper transporting P-type ATPase comprising of the three N-terminal metal binding (M1, M2 and M3), nine transmembrane (numbered 1–9), E1-E2 ATPase and hydrolase domains. **(B)** Amino acid sequence alignment of the putative Leishmanial copper transporting P-type ATPase (accession number: XP_001685970.1) with copper transporting P-type ATPases from *Trypanosoma cruzi* (XP_810568.1), *Pseudomonas putida* (AAP88295.1), *Saccharomyces cerevisiae* (AJU90155.1), *Drosophila melanogaster* (NP_001259466.1), *Danio rerio* (NP_001036185.1), *Mus musculus* (AAA57445.1), *Homo sapiens* (AAA35580.1). Conserved CXXC motifs in the N-terminal domain, actuator (A) domain TGE loop, DKTGT motif in the phosphorylation (P) domain, TGDN and GXGXND and SXHP motifs in the nucleotide (N) domain ATP-binding site are highlighted. Conserved CPC, YNXXXXP and MXXSS motifs are highlighted in the predicted seventh, eighth and ninth transmembrane domains respectively. **(C)** Agarose gel electrophoresis showing amplification of *LmATP7* gene and its transcriptional expression as indicated by 3.5 kb PCR products amplified from both total genomic DNA (left image) and cDNA (reverse transcribed from mRNA) (right image) **(D)** Homology model of N-terminal copper binding domains of LmATP7. (a) Modelled structure of HM1 domain (yellow) with “CRGC” motif shown in inset. (b) Modelled structure of HM2 domain (green) with “CTSC” motif shown in inset. (c) Modelled structure of HM3 domain (blue) with “CASC” motif shown in inset. (d) Modelled structure of Hm2 and HM3 domains combined together (red). (e) Superposition of the individual Hm2 (green) and HM3 (blue) domains on the combined model (red).

### N-terminus Copper binding motifs of LmATP7 share high homology with mammalian Cu-ATPases

PIB-ATPases are heavy-metal-transporting ATPases characterized by the unique presence of Heavy metal (HM) binding sites containing Cys-X-X-Cys motifs at the cytosolic N-terminal domain. Mammalian copper ATPases harbours six such motifs on the amino terminus. This motif sequesters copper and facilitates its transport across the membrane to the lumen [30]. Using homology modelling based on structures of the amino-terminal HM domains of metal ATPases available in the Protein database (PDB), we determine if the novel ATPase that we predicted in the *Leishmania* genome belongs to the family of heavy metal transporting ATPases. We detected three putative HM motifs with the conserved Cys-X-X-Cys motif in LmATP7. The sequence of the three domains that are used for homology modelling are included in the Materials and methods section. Based on E-value of alignment (0.65E-04) the template that was chosen for homology modelling of HM1 was *Bacillus subtilis* CopA (PDB ID: 1OPZ) with the percentage sequence identity between the template and the model normalized by the lengths of the alignment being 30. The modelled structure with the lowest DOPE score of -4818.916016 was selected for further validations and simulation studies (Fig. 2D).

Initially HM2 and HM3 were modelled together since they are in close proximity to one another and hence thought to have evolutionarily co-evolved as compared to HM1 which is sequence-wise farther away. Based on lowest E-value of alignment (0.48E-10), a NMR structure of soluble region of P-type ATPase CopA from *Bacillus subtilis* (PDB ID: 1P6T) was the best template for these two domains combined together. Both the human copper ATPases (ATP7A or ATP7B) lost out on homology modelling both in terms of sequence similarity (29 %) as well as E-value of alignment (0.14E-08). The modelled structure with the lowest DOPE score of -12928.405273 was selected for validation and simulation studies (Fig. 2D).

Surprisingly, when HM2 and HM3 are modelled separately, they show a better homology modelling with human copper ATPases as compared to bacterial ones. For HM2, based on the lowest E value of alignment (0.32E-06) the template which was found suitable for homology modelling was the solution structure of the metal binding domain 4 of human ATP7B (PDB ID: 2ROP) with the percentage sequence identity between the template and the model normalized by the lengths of the alignment being 36. Though there was also a significant percentage sequence identity (45 %) between our model and *Bacillus subtilis* CopA (PDB ID: 1OPZ) and an even lower E value of 0.51E-07, it was rejected based on a huge mismatch in the DOPE per residue score vs alignment position graph between the CopA template and the modelled structure of *Leishmania* HM2 ((Fig. S2). The modelled structure with a lowest DOPE score of -4788.835449 was selected for validations and simulation studies (Fig. 2D).

For HM3, based on the lowest E value of alignment (0.68E-04) the template which was found suitable for homology modelling was the solution structure of the metal binding domains 6 of human ATP7A (PDB ID: 1YJV) with the percentage sequence identity between the template and the model normalized by the lengths of the alignment being 46. Though there was also a significant percentage sequence identity (49 %) between our model and *Bacillus subtilis* CopA (PDB ID: 1OPZ) and an even lower value E-value of 0.32E-05, it was rejected based on a huge mismatch in the DOPE per residue score vs alignment position graph between the CopA template and the modelled structure of *Leishmania* HM3 (Fig. S2). The modelled structure with a lowest DOPE score of -5403.311035 was selected for validations and simulation studies (Fig. 2D).

The above mentioned four models with the lowest DOPE scores obtained through homology modelling were first equilibrated as mentioned in Materials and Methods section followed by a final 1000 ns unrestrained NPT equilibration. The final structures were then obtained by averaging out the last 100 ns of the production runs (Fig. 2D).

### Intracellular localization of LmATP7 in the promastigote and mammalian cells is consistent with other characterized orthologues

The human orthologues ATP7A and ATP7B localize at the trans-Golgi network and traffic in vesicles to the plasma membrane upon copper treatment [31,32]. To determine if LmATP7 localizes in the plasma membrane of *Leishmania*, we cloned and expressed *LmATP7-GFP* in promastigotes (Fig. 3A) and co-labelled it with a *bona fide Leishmania* specific plasma membrane marker GP63 (Fig. 3B). High colocalization of the two proteins was observed; with a small fraction of intracellular puncta of LmATP7-GFP that did not localize with the marker (Fig. 3B,C). LmATP7-GFP staining by anti-GFP was specific and distinct from that of the control parasites that overexpressed only GFP and showed a non-specific signal that was distributed throughout the cell and the wild type parasite that did not show any GFP signal (Fig. 3A). Further to substantiate our findings, FM4-64FX was utilised to determine LmATP7-GFP localization at two time points of the dye uptake, 1 min and 10 min (Fig. 3D,E). FM4-64FX dye labels the plasma membrane at an early time point (1 min) and with time it is endocytosed where it labels the intracellular vesicles (10 min). A high colocalization coefficient was recorded between the marker and transporter at both time points with 10 min time point showing statistically more marker-transporter association than 1 min time point (Fig. 3E). This is suggestive of plasma membrane as well as endocytic vesicular localization of the protein. Upon 2 hours of 50µM copper treatment, at the 10 min time point of FM4-64 uptake, we observed a high persistence of LmATP7 and dye colocalization similar to basal copper condition (Fig. 3F,G), indicating there is no appreciable change in LmATP7 localization upon copper treatment.

**Fig 3.**
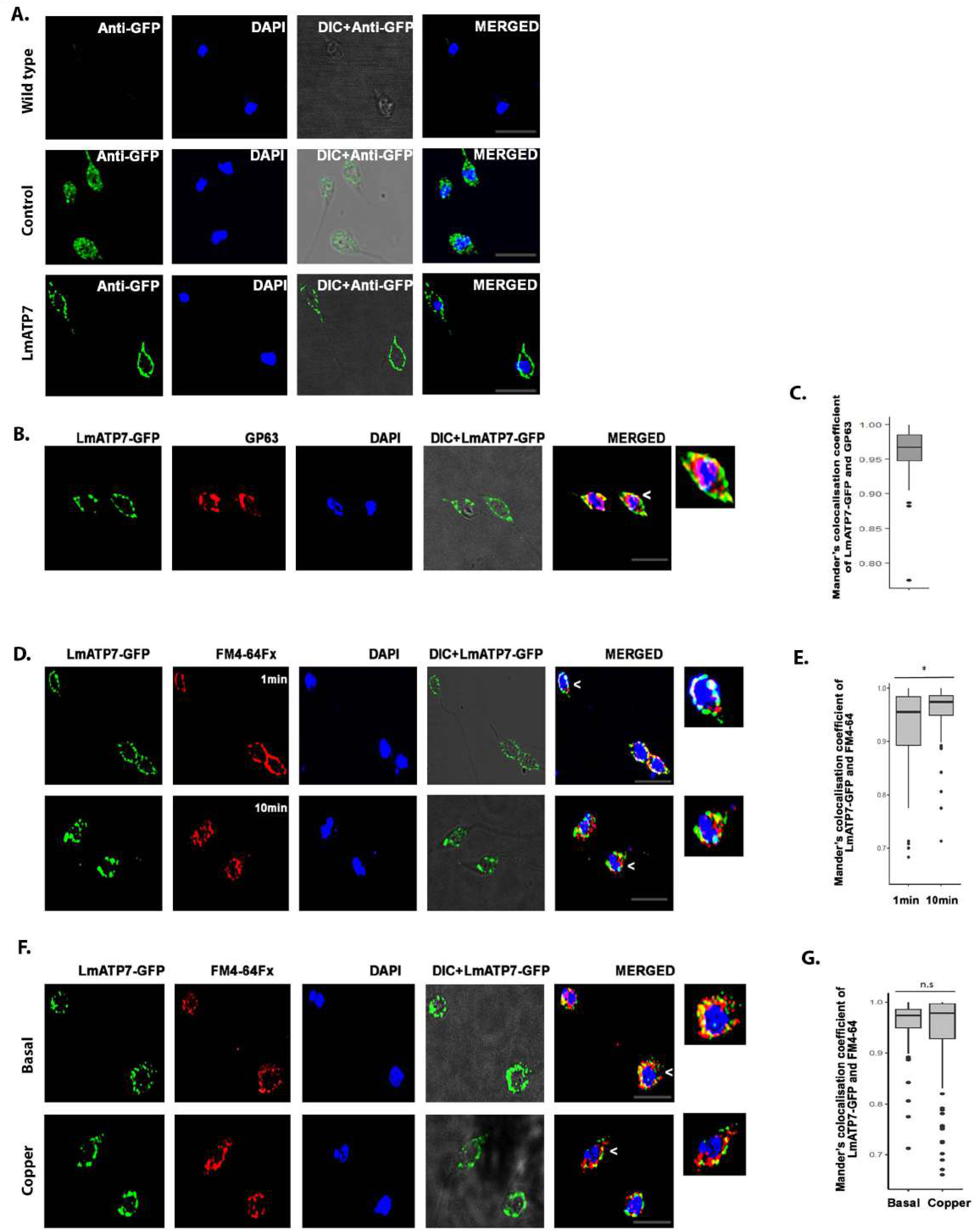
Intracellular localization and expression of LmATP7. **(A)** Wild type, vector control (Control) and *L. major* promastigotes stably expressing LmATP7 as a C-terminal GFP-tagged protein (LmATP7) (right column) were immunostained with antibodies against GFP (green). Promastigote nuclei were stained with DAPI (blue). Scale bar: 5 μm. **(B)** *L. major* cells stably expressing LmATP7–GFP were co-stained for GFP (green) and plasma membrane marker GP63 (red). The merged images show colocalization of proteins (yellow). DAPI (blue) was used to stain the nucleus. The magnified inset corresponds to the region of the merged image marked by arrowhead indicating the association of LmATP7-GFP and GP63. Scale bar: 5 μm. **(C)** Mander’s colocalization coefficient of LmATP7-GFP and GP63 represented by a box plot. Sample size (n) :114. **(D)** *L. major* cells stably expressing LmATP7–GFP were co-stained for GFP (green) and FM4-64FX (red) (1min or 10min incubation). The merged images show colocalization of proteins (yellow). DAPI (blue) was used to stain the nucleus. The magnified inset corresponds to the region of the merged image marked by arrowheads indicating the association of LmATP7-GFP and FM4-64FX. Scale bar: 5 μm. **(E)** Mander’s colocalization coefficient (MCC) of LmATP7-GFP and FM4-64FX represented by a box plot. Sample size (n) for 1min :93, 10min: 102. * indicates significance of difference in the MCC of LmATP7-GFP and FM4- 64FX upon 1 min or 10 min incubation. *P≤0.05 (Wilcoxon rank-sum test). **(F)** *L. major* cells stably expressing LmATP7–GFP under basal and copper treated conditions (50µM for 2 hrs) were co-stained for GFP (green) and FM4-64FX (red) (10 min). The merged images show colocalization of proteins (yellow). DAPI (blue) was used to stain the nucleus. The magnified inset corresponds to the region of the merged image marked by arrowheads indicating the association of LmATP7-GFP and FM4-64FX. Scale bar: 5 μm. **(G)** Mander’s colocalization coefficient of LmATP7-GFP and FM4-64FX represented by a box plot. Sample size (n) for basal:117, copper treated: 113. * indicates significance of difference in the MCC of LmATP7-GFP and FM4-64FX upon basal and copper treated conditions. n.s : non-significant (Wilcoxon rank-sum test).

### LmATP7 is a *bona fide* copper transporter

We hypothesized that if LmATP7 functions as a copper transporter, then parasites overexpressing the protein should have a more efficient copper export mechanism compared to the wild-type counterpart. ICP-OES (Inductively coupled plasma optical emission spectrometry) was performed to determine intracellular copper level of wild type and LmATP7 overexpressing *Leishmania* strain. Higher expression of *LmATP7* was confirmed in promastigote population that stably overexpressed *GFP-LmATP7* (Fig. S4). Copper level was approximately three fold less in the LmATP7 overexpressed strain as compared to the wild type parasite (Fig. 4A) suggestive of the role of LmATP7 in transporting copper out of the cell.

**Fig 4.**
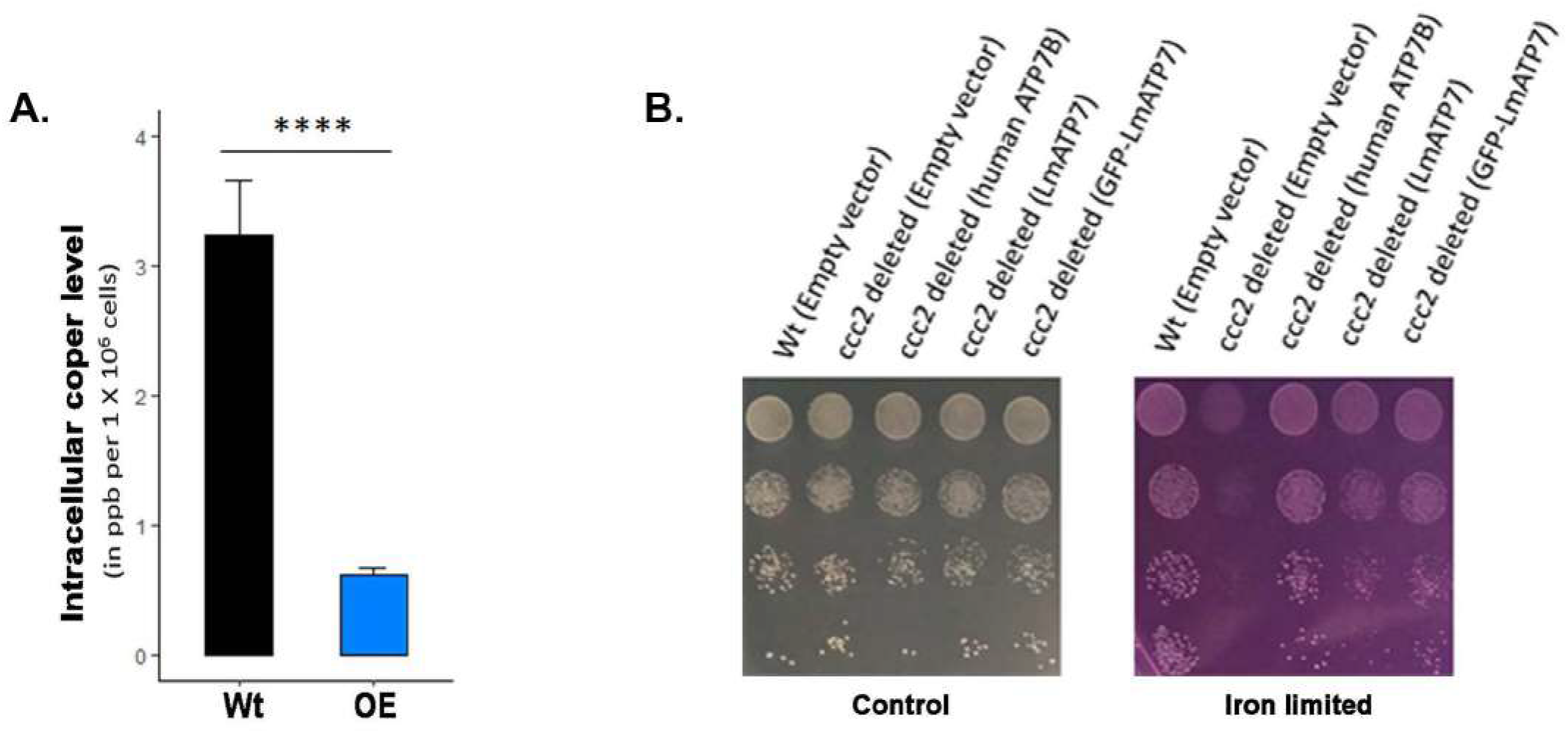
LmATP7 transports copper. **(A)** Bar diagram showing the intracellular copper content (in ppb) of 1X10^6^ cells of wild type (Wt) and LmATP7 overexpressing (OE) *L. major* promastigotes as measured by ICP- OES. Error bars represent mean ± SD of values from three independent experiments. * represents the significance of difference in the amount of internal copper content of wild type and LmATP7 overexpressing *L. major* promastigotes. ****P≤0.0001 (Student’s t-test) **(B)** Complementation of ccc2 mutant yeast by LmATP7. Wild-type (Wt) and deletion (ccc2) yeast strains were transformed with constructs mentioned within the parentheses. Dilution spotting was performed on SD-Ura plate (Control) and 250 µM ferrozine treated SD-Ura plate (Iron limited) with yeast transformants (OD_600_ = 0.2, 0.02, 0.002, 0.0002). Images of the plates were taken after 3 days of incubation at 30°C.

To further determine if LmATP7 is a *bona fide* copper ATPase, we used a functional assay that has been developed for human P-type Cu-ATPases, ATP7A/B and its mutants. This assay is based on functional complementation of the CCC2 deletion mutant (*ccc2Δ*) of the yeast *Saccharomyces cerevisiae* [33]. Ccc2p, the P-type ATPase copper transporter of *S. cerevisiae* transports copper to the plasma membrane oxidase Fet3p, which then works with a high affinity iron importer Ftr1p to import iron from the medium. In the yeast deletion mutant *ccc2Δ,* copper is not incorporated into Fet3p leading to deficiency in high affinity iron uptake. In iron-sufficient medium, the *ccc2Δ* can grow well owing to the ability of low affinity iron uptake systems to supply iron to the cells. However, in iron limited medium, this mutant fails to grow as Fet3p-Ftr1p, which becomes essential under these conditions, is non-functional. ATP7B, in previous studies, has been shown to complement the *ccc2Δ* mutation allowing the mutant cells to grow in iron-limited medium, establishing this as a convenient functional assay for P-type Cu-ATPases.

The *S*.*cerevisiae ccc2Δ* mutant was accordingly transformed with either *LmATP7* and *GFP-LmATP7* expressed downstream of the strong TEF promoter in a single copy centromeric vector (p416TEF-*LmATP7,* p416TEF*-GFPLmATP7*) or the empty vector pTEF416 or the human *ATP7B* that was similarly expressed as a positive control (p416TEF-*ATP7B*). The transformants that appeared were evaluated for growth in iron deficient and iron-sufficient (control) media by serial dilution assays. We observed that expression of LmATP7 and GFP-LmATP7 in *ccc2Δ* was able to complement the mutation restoring their growth in iron-limited medium. In contrast, the cells bearing the vector control failed to grow. The p416TEF-*ATP7B* construct complemented more strongly as compared to LmATP7 and GFP-LmATP7 and may reflect either differences in expression of the two genes in yeast, or possible differences in functional efficiency. Importantly, LmATP7 showed complementation indicating that this leishmanial protein was able to functionally complement *CCC2* in vivo (Fig. 4B). Hence *LmATP7* can be functionally ascertained as the IB P-type copper transporting ATPase in *L. major.* Since GFP-tagged LmATP7 has been used in other cell based experiments in this study, we ensured that the GFP chimera can complement *CCC2* indicating that GFP did not interfere with the correct LmATP7 localization on which its functioning depends.

### LmATP7 imparts copper tolerance to *L. major* promastigotes

Since we have ectopically expressed and established LmATP7 as a Cu-ATPase in yeast, we now investigated if the protein is capable of regulating copper homeostasis in *Leishmania* as copper treatment has been shown to upregulate ATP7A in mammalian cells [34]. For this, we attempted to generate both overexpressing and knockout *L. major* lines of this gene. Although we could successfully generate GFP-tagged LmATP7 overexpressing line and heterozygous mutant strain of LmATP7 using pXG-NEO, our repeated attempts failed to develop the complete knockout of this gene in *L. major*. Together it strongly indicated that LmATP7 plays a crucial role in parasite survivability. To determine if copper has a role on LmATP7 regulation in *Leishmania*, we challenged wild type (Wt), *LmATP7* overexpressing (LmATP7-OE) and *LmATP7* heterozygous deletion (LmATP7-Het) *L. major* promastigotes with 50µM copper for 2 hrs or not and determined the transcript levels. Interestingly, in all cases we observed increase in the transcript level while the cells were treated with copper indicating the possible role of LmATP7 in maintaining *Leishmania* copper homeostasis (Fig. 5A). Further to verify if this gene imparts any resistance towards Cu mediated toxicity on the parasites we further grown wild type, vector control (GFP-only), LmATP7-OE or LmATP7-Het promastigotes in presence of 0- 500 µM of external Cu and systematically observed the parasite growth kinetics at 0- 72 hrs post incubation. As evident from Fig. 5B-D both wild type and vector control (Fig.S5) *L. major* strains showed reduced growth with increasing copper in the medium starting from 24 hrs post incubation indicating cytotoxic effect of excess copper. In fact, at 500 µM concentration of external copper, wild type and vector control promastigotes count were lower than the initial seeded number at 24 hrs post incubation indicating cell death. Interestingly, LmATP7 overexpressing promastigote count was significantly higher than the controls at such heightened copper level throughout the time points which could even tolerate 500 µM of external Cu stress. Although its growth decreased gradually with increasing copper conditions, it consistently maintained a higher growth rate than the controls. This indicates an enhanced copper export activity in LmATP7 overexpressed promastigotes (also evident from Fig. 4A), allowing a relatively better internal copper regulation and reduced cytotoxic effects. Importantly, LmATP7-Het cells struggled to grow at a comparable rate with wild type cells even in absence of external Cu. Also, starting from 50 µM of Cu treatment the cell count decreased gradually with increasing Cu concentration as well as with time. We have also shown the parasite growth between wild type, LmATP7-OE and LmATP7-Het cells at 72 hrs post incubation in presence of 0- 500 µM of Cu concentration and illustrated our findings in a lucid way (Fig. 5E, bottom right). Both the controls were also able to survive and maintain their growth to some extent under excess external copper, probably owing to their endogenously expressed LmATP7. At extremely high copper, endogenous LmATP7 seemed to be insufficient to maintain internal copper homeostasis, leading to tremendous growth retardation and cell death. The results provided the first evidence of the role of the newly characterized leishmanial Cu-ATPase, LmATP7, in copper tolerance.

**Fig 5.**
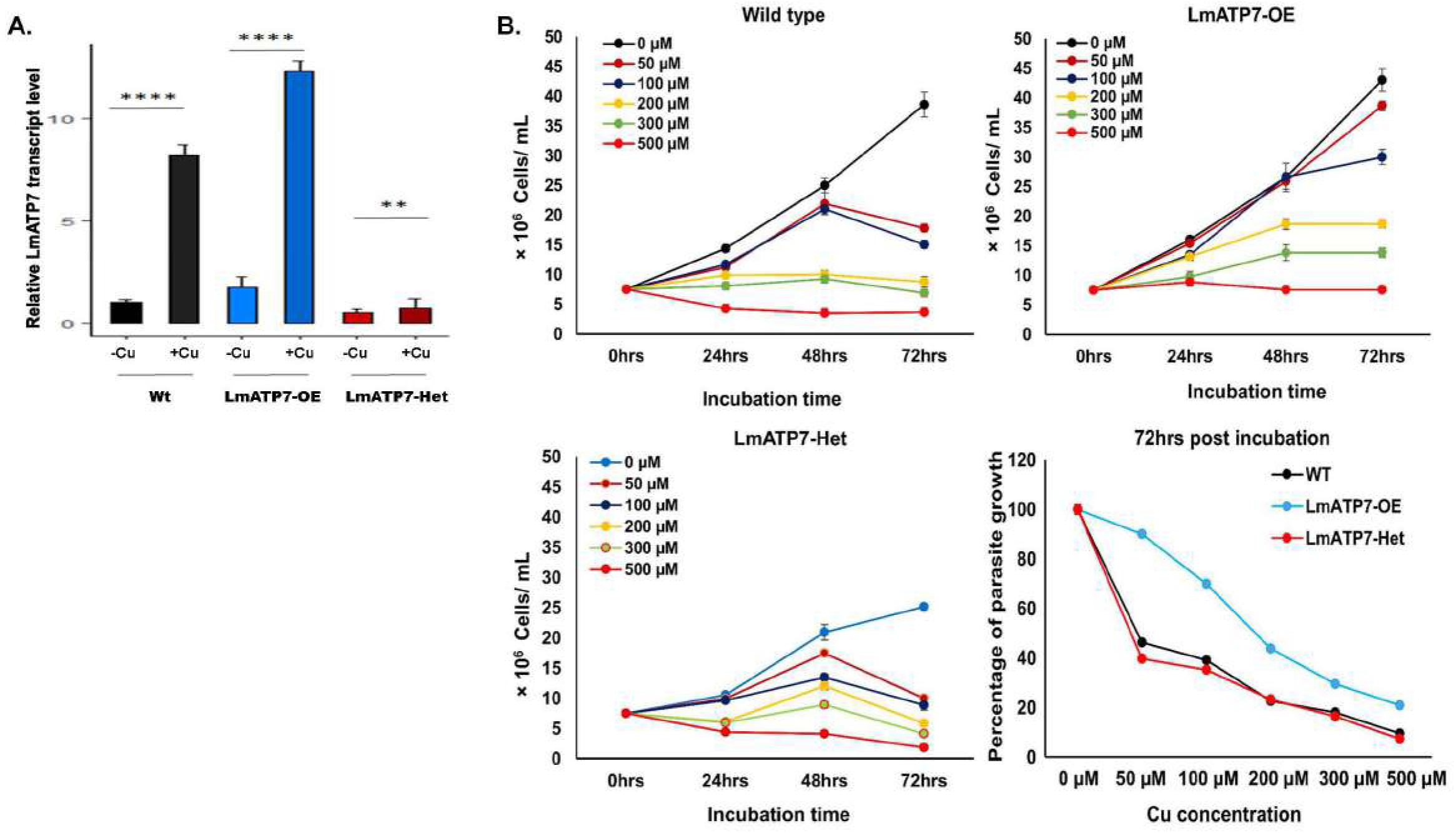
LmATP7 imparts copper tolerance to *L. major* promastigotes. **(A)** Bar diagram showing LmATP7 transcript level in wild type (Wt), LmATP7 overexpressing (LmATP7-OE) and LmATP7 heterozygous deletion (LmATP7-Het) strains of *L. major* grown either normally or in presence of 50µM of external copper for 2 hrs. Expression of LmATP7 was normalized against rRNA45 gene. Error bars represent mean ± SD of values calculated from three independent experiments. * indicate significance of difference in expression of LmATP7 upon copper treated and untreated conditions. **P≤0.01, ****P≤0.0001 (Student’s t-test). **(B)** Wild type, LmATP7 overexpressing (LmATP7-OE) **(C)** and LmATP7 heterozygous deletion (LmATP7-Het) **(D)** *L. major* promastigotes were grown at 0- 500 µM externally added copper for 24 hrs, 48 hrs and 72 hrs. Initially, 7.5×10^6^ cells were added to the media at time point 0 hrs and further at the above mentioned time-points, cells were counted by haemocytometer-based cell counting. Error bars represent mean ± SD of values from three independent experiments. **(E)** Line graph showing the comparison between growth of Wild type (WT), LmATP7 overexpressing (LmATP7-OE) and LmATP7 heterozygous deletion (LmATP7-Het) promastigotes in presence of the indicated concentrations of external copper (0- 500µM) at 72 hrs post incubation. Error bars represent mean ± SD of values from three independent experiments.

### Macrophages channelize copper to intracellular compartments harbouring *Leishmania* amastigotes

To further delineate the role of LmATP7 in a physiological condition, we explored host-*Leishmania* interaction with respect to copper stress subjected by the host upon parasite and the parasite’s reciprocating response thereon to evade the stress. At the outset, we examined alterations of macrophage transcripts levels (if any) of the key genes *(ATP7A, CTR1, CTR2, ATOX1)* of copper homeostatic pathway during *Leishmania* infection. ATP7A is a P-type ATPase that provides copper to copper dependent enzymes. When copper is in excess, ATP7A which is a Golgi resident protein, acts as an exporter and traffics out of Golgi in vesicles to remove the copper from the cell [35]. CTR1 and CTR2 are copper importers located in plasma membrane and endo-lysosomal membranes respectively while Atox1 is a cytosolic copper chaperone that carries copper from CTR1 to ATP7A [36,37]. J774A.1 macrophages were infected with *L. major* promastigotes and infection was confirmed and measured by counting the nuclei of amastigotes in the macrophage. Transcripts of Cu-ATPase, *ATP7A*, copper transporters, *CTR1*, *CTR2* and copper chaperone, *ATOX1* were measured 12 hrs and 30 hrs post infection. Interestingly, all four genes were downregulated at the shorter post infection time-point; however, at 30 hrs post infection, *ATP7A* exhibited ∼1.8X upregulation. Other transcripts also exhibited modest upregulation (Fig. 6A). The results suggested that host copper uptake and utilization is downregulated during early stages of infection. However, the host exerts its copper mediated response at a later stage with upregulation of *ATP7A*, *CTR1* and *CTR2*.

**Fig 6.**
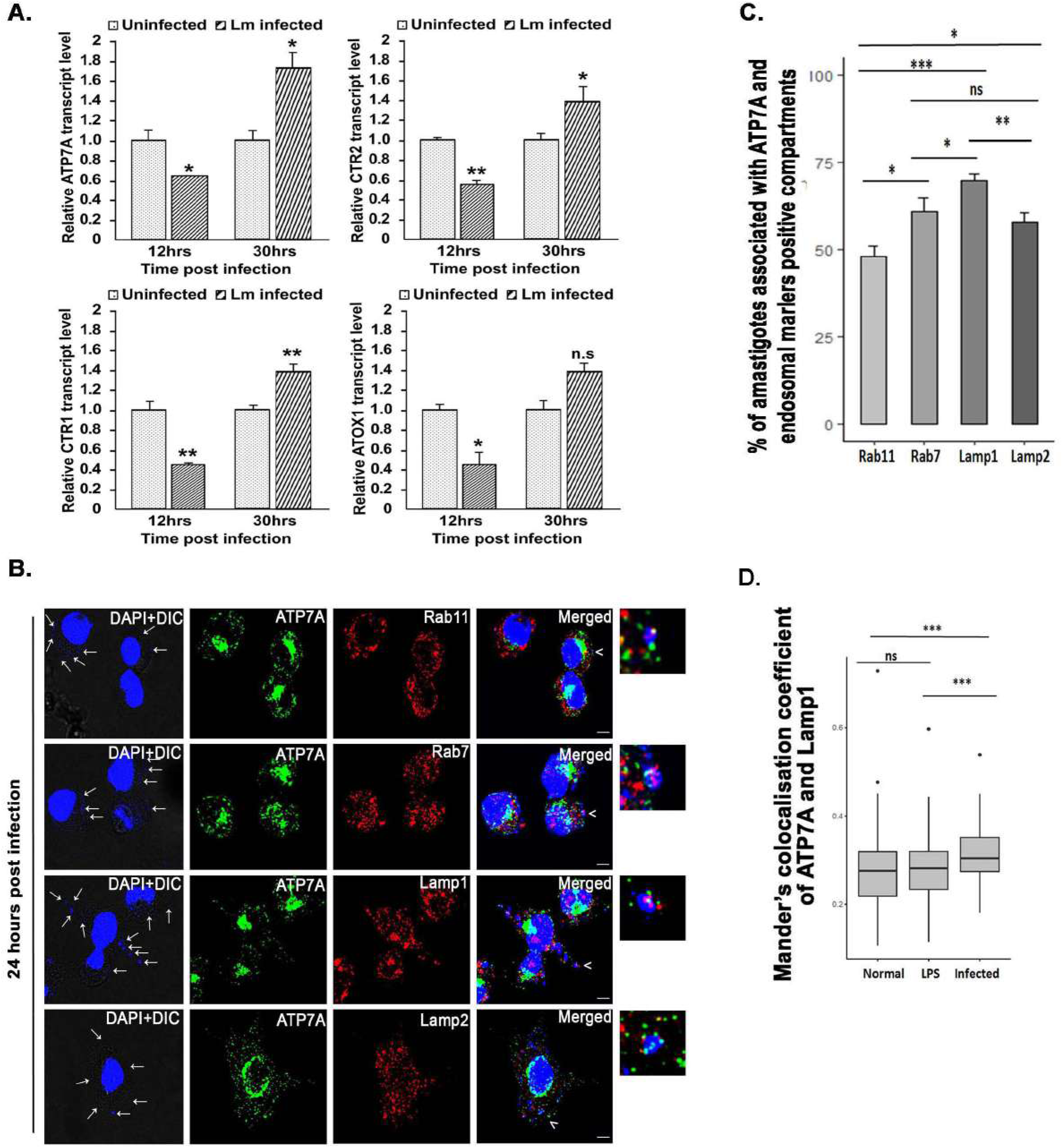
*Leishmania* infection elicits a host response that is mediated through its copper homeostasis genes. **(A)** J774A.1 macrophage cells were either infected or not with *L. major* promastigotes for 12 hrs and 30 hrs. Bar diagram shows transcript levels of macrophage copper homeostasis genes *ATP7A, CTR1, CTR2*, and *ATOX1* normalized against *β-Actin* mRNA levels. Error bars represent mean ± SD of values calculated from three independent experiments. * indicates significance of difference in the transcript level of the genes between infected macrophages with respect to those in uninfected macrophages at the particular time points, (12 hrs, 30 hrs). *P≤0.05, **P≤0.01, n.s : non-significant (Student’s t-test). **(B)** J774A.1 macrophages were infected with *L. major* promastigotes and stained with anti-ATP7A (green) and anti-Rab11 (red) or anti-Rab7 (red) or anti-Lamp1 (red) or anti-Lamp2 (red) at 24 hrs post infection. The merged images represent colocalization of ATP7A with *Leishmania* amastigote nuclei (blue) and endosomal markers (Rab11, Rab7, Lamp1, Lamp2) (red). Both macrophage and *Leishmania* nuclei were stained with DAPI (blue). White arrows indicate intracellular parasites in infected cells (smaller nuclei). The magnified inset corresponds to the region of the merged image marked by arrowheads indicating the association of ATP7A and endosomal markers with *Leishmania* positive endosomes. Scale bar: 5µm. **(C)** Bar plot represents the percentage of *L. major* amastigote nuclei associated with both ATP7A and the indicated endosomal markers (Rab11, Rab7, Lamp1, Lamp2) of J774A.1 macrophages. The number of *Leishmania* nuclei counted to obtain the data for each condition are 197, 189, 274, 243 (for Rab11, Rab7, Lamp1, Lamp2 respectively) from three independent experiments. Error bars represent mean ± SD of values calculated from three independent experiments. * indicates significance of difference in the percentage of amastigote nuclei associated with both ATP7A and the indicated endosomal markers. *P≤0.05, **P≤0.01, ***P≤0.001, n.s : non-significant (Wilcoxon rank-sum test). **(D)** Mander’s colocalization coefficient between ATP7A and Lamp1 from uninfected (Normal), LPS treated (LPS) and *L. major* infected J774A.1 macrophages (Infected) represented by a box plot. Sample size (n) for normal: 83, LPS treated: 79, Infected: 75. * indicates significance of difference in the colocalization of ATP7A and Lamp1 in the three different samples. ***P≤0.001, n.s: non-significant (Wilcoxon rank-sum test).

To export excess copper, ATP7A traffics to vesicle out of TGN (Trans-Golgi network) when cells are treated with high copper [32,38,39]. Macrophage ATP7A is capable of trafficking upon copper treatment (Fig.S3) Hence, as a function of copper transport, we probed intracellular localization and TGN exit of ATP7A at 24 hrs post-*Leishmania* infection. The time point fits between the heightened parasite response and heightened host response. Interestingly, ATP7A traffics out of Golgi in vesicles in *L. major* infected macrophages (Fig. 6B). ATP7A in uninfected macrophages remained clustered at the Golgi (Fig. S3). Since *Leishmania,* post-internalization, localizes in the macrophages throughout the endosomal pathway, we determined the nature of the ATP7A positive compartment that harbours this amastigote parasite. To scan the entire endocytic pathway we measured percentage of amastigote nuclei associated with both ATP7A and the endosomal/endo-lysosomal markers, Rab11 (recycling endosome), Rab7 (late endosome) and Lamp1/2 (endo-lysosome). We noticed that colocalized ATP7A-amastigote nuclei distributes throughout the endosomal pathway (Fig. 6B,C); with highest localization at the Lamp1 positive compartments (Fig. 6C, D).

### LmATP7 is important for the intracellular survival and infectivity of *L. major*

It is now obvious that as a host response, the macrophage deploys the copper transport machinery to exert stress on the intracellular parasites. We argue that *Leishmania* LmATP7 bypasses copper stress efficiently brought-upon by the host macrophage during their intracellular life cycle. Stable lines of *L. major* expressing GFP-LmATP7 or heterozygous deletion mutant LmATP7-Het generated as mentioned earlier were used subsequently to infect J774A.1 macrophages under normal and increased copper conditions. Empty vector (pXG-GFP+) expressing and wild type promastigotes were used as controls. Corroborating with our hypothesis, we found that *L. major* amastigotes overexpressing LmATP7 were significantly higher in abundance as compared to both the controls at 24 hrs post infection (Fig. 7A). This increase was consistent both at normal and copper-treated conditions suggesting when overexpressed, LmATP7 by exporting copper from *Leishmania*, provided increased survivability to the amastigotes (Fig. 7B, S6A). Previous studies showed bactericidal activity is promoted by intracellular bio-available copper in macrophages which was further facilitated by external copper treatment [23]. We also observed a similar phenotype of reduced intracellular amastigotes with increasing copper level. As illustrated in Fig. 6B, under physiological copper condition, activated macrophages increase copper import and partially channel copper via ATP7A towards the endo-lysosomal compartments. But LmATP7 overexpressed protein was probably more efficient in removing excess copper from the *Leishmania* cells as compared to the endogenously expressed one resulting in higher amastigote per macrophage cell. However, under the copper treated condition LmATP7-Het failed to establish a successful infection. As compared to wild type control, even in absence of Cu the parasite burden is more than 2 folds low in case of LmATP7-Het. In fact, the trend remained same with increasing concentration of Cu treatment (Fig. 7A, B and S6A). Interestingly, we also wanted to check whether this gene plays any role in parasite infectivity as well as intracellular replication of amastigotes. As shown in Fig. S6B there was ∼20% increase in percent of infected macrophages while the J774A.1 cells were infected with LmATP7-OE at both 24 and 48 hrs post infection as compared to its vector control and wild type counterpart. But the LmATP7-Het cells showed a significant ∼25% decrease in percent of infected macrophages at 24 hrs post infection as compared to wild type *L. major*. In fact, at 48 hrs post infection there was ∼30% decrease (Fig. S6B). Similar to this observation when we extended our infection study with all those four strains upto 48 hrs in absence of Cu we have seen that there was about 2 folds increase in parasite burden at 48 hrs post infection with LmATP7-OE as compared to wild type or vector control. However, at that time point while the wild type *L. major* could maintain its population successfully, LmATP7-Het cells showed about 3 folds decrease with the parasite burden of ∼0.7, i.e., less than one amastigotes/ macrophage cell (Fig. S6C). Overall, the experimental data revealed the importance of LmATP7 in neutralizing toxic levels of copper from the host on *Leishmania* amastigotes. Also, collectively our study establishes that LmATP7 is crucial for both the survivability as well as infectivity of *L. major* parasite.

**Fig 7.**
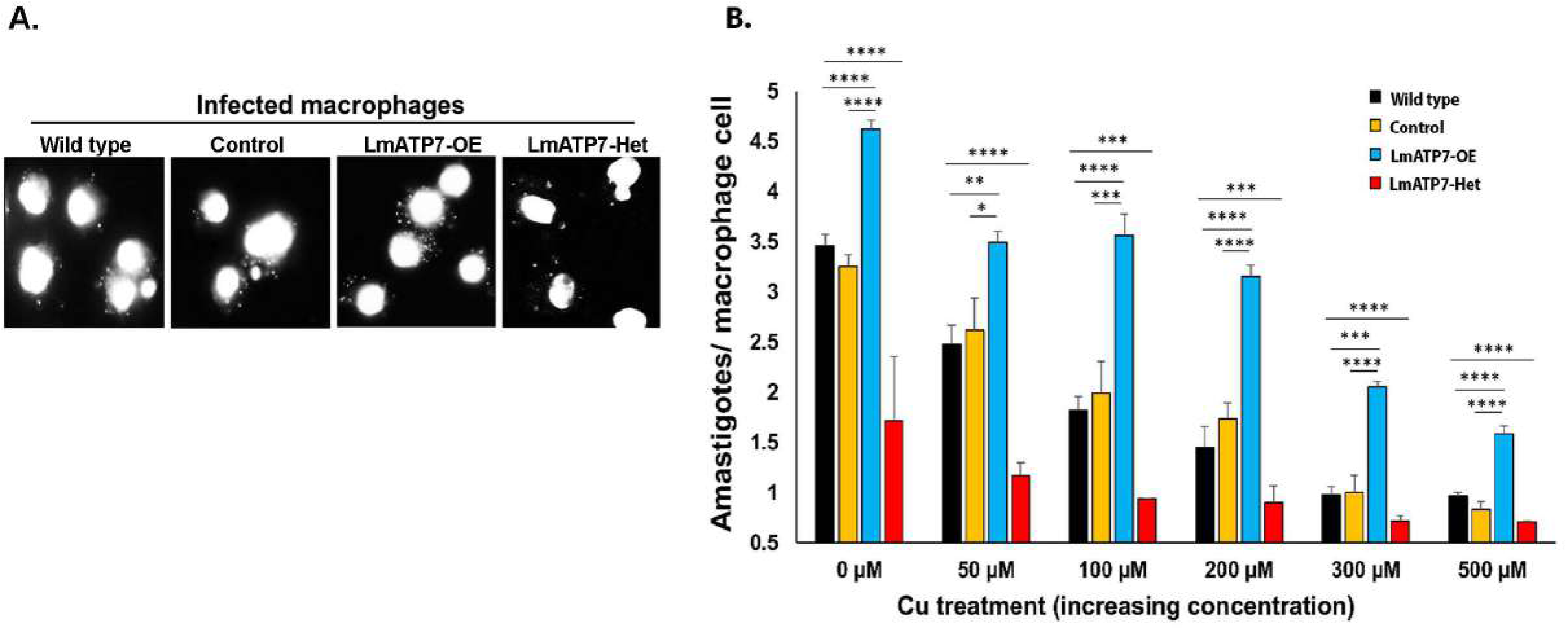
LmATP7 is key for amastigote survivability in macrophages in basal and elevated copper. **(A)** Representative DAPI stained epifluorescence microscopy images of three independent experiments showing J774A.1 macrophages infected with Wild type, vector control (Control), LmATP7 overexpressed (LmATP7- OE) or LmATP7 heterozygous deletion (LmATP7-Het) *L. major* strains post 24 hrs of infection. Larger nuclei represent macrophage cells surrounded by smaller amastigote nuclei of the parasite. **(B)** Graphical representation of amastigotes/ macrophage count at 24 hrs post infection. J774A.1 macrophages were infected with Wild type (black bar), vector control (Control, yellow bar), LmATP7 overexpressed (LmATP7-OE, blue bar) or LmATP7 heterozygous deletion (LmATP7-Het, red bar) strains in presence of indicated concentrations of copper (0- 500µM). At least 100 macrophages were counted from triplicate experiments. Error bars represent mean ± SD of values from three independent experiments. * indicates the significance of difference in amastigote/ macrophage count in case of Control, LmATP7-OE and LmATP7-Het strains compared to Wild type cells at a particular copper concentration. *P≤0.05, **P≤0.01, ***P≤0.001, ****P≤0.0001; (Student’s t-test).

**Fig 8.**
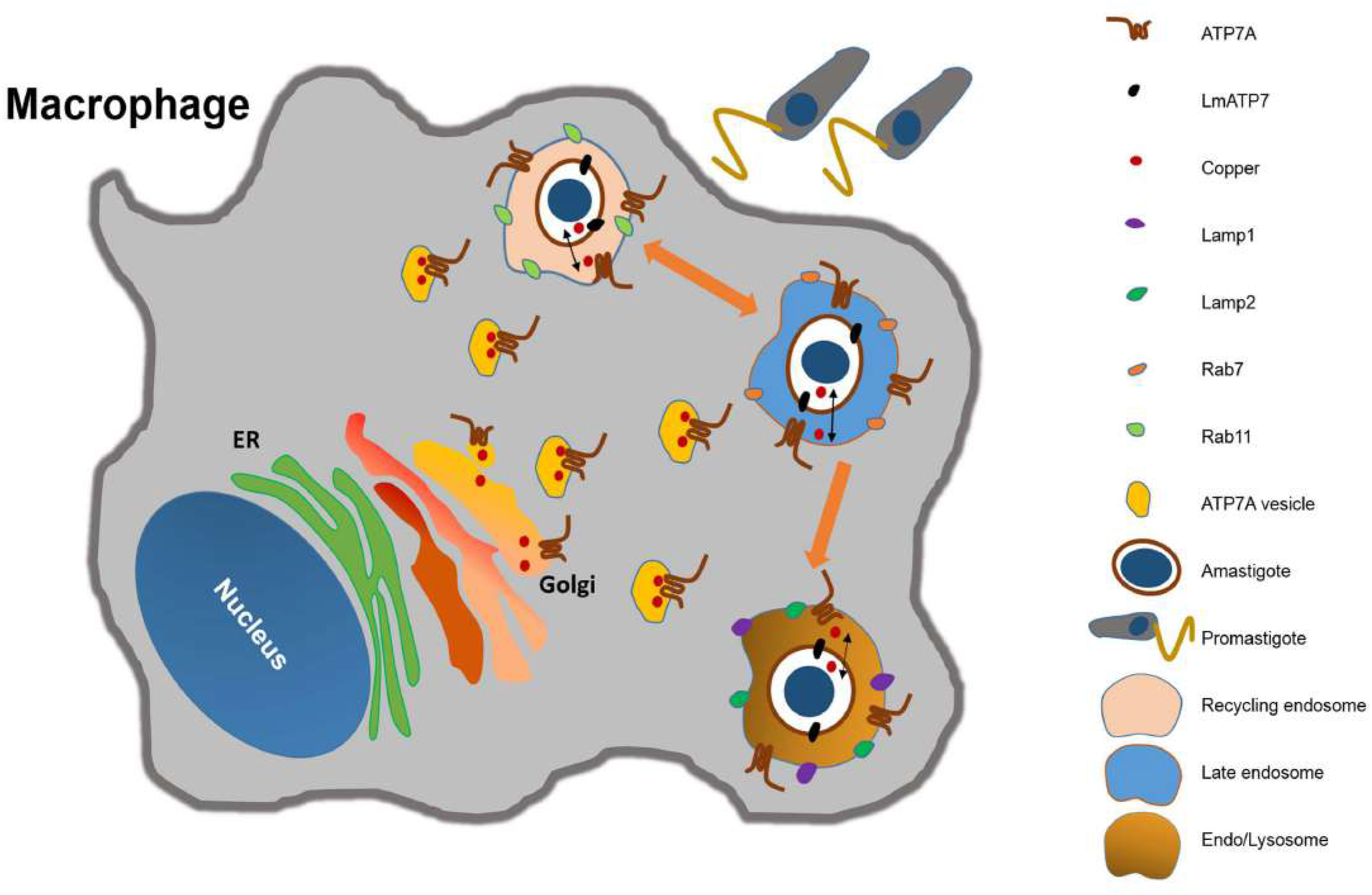
Proposed model describing the role of parasite and host copper P-type ATPases in *Leishmania* infection. Macrophages channelize copper via ATP7A trafficking from Golgi to the *Leishmania* positive endo/lysosomal compartments. *Leishmania* amastigotes, in turn, export copper via LmATP7 to reduce the host-mediated copper stress in order to thrive within the host as indicated by black double headed arrows. Orange arrows indicate the endo-lysosomal pathway followed by amastigote after its entry into the macrophage.

## Discussion

Host-parasite interaction is a rapidly evolving field. Host-parasite interaction has served as an apt example of ‘red queen hypothesis of winner less antagonistic coevolution’ [40]. Host and the parasite co-evolves where parasites adapt to neutralize any response that is employed by the host to evade parasite invasion. Macrophage cells, being the sentinel of our immune system play pivotal role in eliminating different pathogenic infections [41]. However, *Leishmania* parasites specifically infect and survive within these cells. Macrophages have been shown to express the key copper-homeostatic genes. Further, it has been demonstrated that in the case of pathogenic yeast, *Candida* sp. infection, macrophage deploys the copper ATPase, ATP7A to exert copper-mediated toxicity over the pathogen [42]. However, there is no such report which addresses if macrophage being the host to *Leishmania* tries to combat the infection using similar mechanism and if so, then how the parasite escapes such host-induced stress. In this context, we have identified and characterized the Cu-ATPase, LmATP7 for the first time in the kinetoplastid parasite genus, *Leishmania*. Furthermore, using the well-established macrophage-*Leishmania* infection model system we found that there is a reciprocal interaction between the copper ATPases of the host and the parasite that determines the fate of parasite survivability and well-being of the antagonistic pair.

We observed that upon increasing concentration of copper treatment wild-type *L. major* promastigotes could overcome the toxic effect of this metal ion. This observation is particularly interesting since the other known bacterial pathogens are reported to be susceptible towards copper concentration as low as 20 µM [23]. Although at 50 µM and higher external Cu treatment, there was a drop in parasite count, *Leishmania* could maintain its population appreciably at a very high Cu concentration of 500 µM. We noticed a similar survival trend even in the intracellular amastigotes in these copper conditions. These two exciting observations collectively suggest that there must be an efficient copper homeostasis mechanism within *Leishmania* which is expressed both in its extracellular promastigote form as well as in its infectious amastigote stage.

Importance of copper in kinetoplastid parasites is unknown. Though the Cytochrome C oxidase complex (that is known to bind copper) has been identified in *Leishmania* and its component Ldp27 has been shown to be crucial in amastigote survival, no information is available about role of copper in the secretory pathway [43]. Additionally, presence of a Cu/Zn-type superoxide dismutase in *Leishmania* glycosomes may indicate the role of a Copper P-type ATPase in incorporating copper to TGN and intracellular organelles [44]. Although the role of iron has been extensively studied in kinetoplastid parasites; there has been limited efforts to understand the role of copper in their survival or pathogenicity. Mammalian Copper ATPase, ATP7A carries out two primary functions. Firstly, it transports copper to enzymes that require the metal as a co-factor for maturation, e.g., Dopamine-β-hydroxylase and tyrosinase. Secondly, upon copper excess in the cell, ATP7A traffics in vesicles to export out the excess copper [32]. In a polarized epithelial cell, as in intestinal enterocytes, ATP7A transports copper through the basolateral surface (blood-side) that is further picked-up in the serum by copper-sequestering proteins, e.g., albumin. However, in unpolarised cells, e.g., macrophages and fibroblasts, the copper transporter vascularizes upon excess copper and localizes at the lysosomes and exports copper by triggering the lysosomal exocytosis pathway [39,45,46]. Thus, it would be interesting to determine if *Leishmania* copper ATPase carries out the dual function of copper detoxification as well as copper transport to the secretory pathway as done by its mammalian counterparts, ATP7A and ATP7B.

Recently it has been demonstrated that *Trypanosome* genome codes for a copper-ATPase [29]. We identified a putative *Leishmania* specific P-type ATPase with three predicted N-terminal copper binding motifs (sharing high homology with mammalian CuATPases) and nine transmembrane domains. It is worth mentioning that lower order eukaryotes carry a single copper-ATPase; however with the advent of cellular apico-basal polarity and increasing tissue complexity in higher organisms, this copper-ATPase branched out into two homologues ATP7A and ATP7B. Our in depth bioinformatics analysis depicted this putative ATPase as a copper transporter which was further confirmed with the yeast complementation study. Like its other protozoan counterpart, *Plasmodium berghei* and *Toxoplasma gondii* [47], the LmATP7 is present both in endosomal compartments as well as in plasma membrane. It is well-known that mammalian CuATPases, in response to high levels of copper, relocalize to the vesicles that eventually fuse with the plasma membrane to remove excess copper from the cell [32,38]. ICP-OES data showing reduced internal copper level in LmATP7 overexpressed *L.major* suggests the role of LmATP7 as a copper exporter. Greater survivability of LmATP7-OE strain and reduced survivability of LmATP7-Het strain with respect to wild type promastigotes when challenged with external copper revealed the importance of LmATP7 in copper tolerance. Moreover its expression is regulated by copper as observed from the qRT-PCR data indicating its role in copper homeostasis mechanism.

To inflict a copper induced stress on *Leishmania*, ATP7A of the macrophage traffics to transport copper to compartments that harbour the parasites. Amastigotes overexpressing LmATP7 exhibit a significantly higher colonization in the macrophages as compared to the wild-type control, further establishing the role of LmATP7 in combating host copper stress. A similar trend of higher parasite survivability of the LmATP7-overexpressing strain was observed upon increasing copper treatment of the macrophages harbouring the amastigotes. It is worth mentioning that at least up to 200 µM Cu treatment the parasite burden in LmATP7 overexpression strain infected macrophages was ≥ 3 (amastigotes/ macrophage cell) which is about two fold higher than the parasite burden in either wild-type *Leishmania* or vector control infected macrophages. Moreover, LmATP7-Het strain exhibited a significantly reduced colonization in the macrophages as compared to the wild type control with and without external Cu treatment. Along with reduced survivability, its infectivity also went drastically down. These results along with repeated failures to generate a complete knockout line for the gene with could suggest its essential nature.

Although it has been reported that several intracellular pathogens try to arrest the phagosome maturation process in the host macrophage cells to avoid transport to lysosomes, *Leishmania* is exceptional since it prefers to reside within the endolysosomal and the phagolysosomal compartment [5,48]. There are numerous reports that demonstrate macrophage cells can actively accumulate and compartmentalize copper in response to developing potential anti-microbial activities [23,49]. In fact, it has been shown that copper concentration within phagolysosomal compartment of macrophages increases significantly when infected with *Mycobacteria* [22]. This might be a crucial step towards parasite killing, since under the acidic environment of the phagolysosome, copper can react with the readily available reactive oxygen and nitrogen species to exert its toxicity [23,50]. Hence, it will be interesting to determine if macrophage cells also utilize copper in a similar fashion to kill *Leishmania* and the reciprocal response of the parasite, if any. Along this line, our observation of parasite mediated downregulation of macrophage *ATP7A*, *CTR1*, *CTR2* as well as *ATOX1* at early time point, i.e., 12 hrs post infection with *L. major* implicates the same for the first time. In the putative series of events, we argue that as a host response, the macrophage upregulates ATP7A followed by its copper transport to the endosomal pathway harbouring the parasite. Subsequently, the parasite responds by upregulating the copper ATPase, LmATP7 to alleviate the host-induced copper stress and eventually survives inside the macrophage.

Recently, it has been shown that *Leishmania* hijacks host regulatory pathways that control expression of a phagolysosomal iron transporter so that it can effectively acquire iron for its survival by increasing the concentration of this metal ion within phagolysosomes [51]. This is certainly a parasite mediated remodelling of host machineries. However, our study shows that unlike in iron homeostasis, host macrophages tries to concentrate copper within *Leishmania* containing phagolysosomal compartment to kill the pathogen. To counteract this toxic effect the pathogen has armoured themselves with the Cu-ATPase, LmATP7. Hence, it will be extremely important to identify the underlying mechanisms behind the parasite mediated downregulation of host’s Cu utilization proteins as well as how LmATP7 could be targeted to develop novel anti-leishmanial drug.

## MATERIALS AND METHODS

Reagents were purchased from Sigma-Aldrich (St. Louis, Missouri, United States) unless mentioned specifically. Primers were obtained from GCC biotech (West Bengal, India) and their sequence details are provided in Table S1.

### Plasmids and antibodies

*Leishmania* expression vector (pXG-GFP+) was a generous gift from Dr. Stephen M. Beverley (Washington University Medical School, St. Louis). Mammalian expression vector (pcDNA3.1+) was a kind gift from Dr. Partho Sarothi Ray (IISER Kolkata).

Following are the antibodies used for experiments: rabbit anti- GFP (#BB-AB0065, Biobharati, Kolkata, India), mouse anti-GP63 (#MA1-81830, Invitrogen, Carlsbad, California, United States), rabbit anti- ATP7A (ab 125137, Abcam, Cambridge, United Kingdom), goat anti- Rab11 (# sc- 6565, Santa Cruz Biotechnology, Dallas, Texas, United States), mouse anti-Rab7 (#sc-376362, Santa Cruz Biotechnology, Dallas, Texas, United States), mouse anti-Lamp1 (DSHB: #H4A3, Iowa City, IA, United States), mouse anti-Lamp2 (DSHB: #H4B4, Iowa City, IA, United States), donkey anti-rabbit IgG (H+L) Alexa Fluor 488 (#A-21206, Invitrogen, Carlsbad, California, United States), donkey anti-goat IgG (H+L) Alexa Fluor 568 (#A-11057, Invitrogen, Carlsbad, California, United States), donkey anti-mouse IgG (H+L) Alexa Fluor 568 (#A10037, Invitrogen, Carlsbad, California, United States). FM4-64FX dye (F34653, Invitrogen, Carlsbad, California, United States) was a generous gift from Dr. Bidisha Sinha (IISER Kolkata).

### Parasite and mammalian cell culture

The *L. major* strain 5ASKH was a kind gift of Dr. Subrata Adak (IICB, Kolkata). *L. major* promastigotes were cultured in M199 medium (Gibco, Thermo Fisher Scientific, Waltham, Massachusetts, United States) supplemented with 15% heat-inactivated fetal bovine serum (FBS, Gibco, Thermo Fisher Scientific, Waltham, Massachusetts, United States), 23.5 mM HEPES, 0.2 mM adenine, 150 µg/ml folic acid, 10 µg/ml hemin, 120 U/ml penicillin, 120 µg/ml streptomycin, and 60 µg/ml gentamicin at pH 7.2 and the temperature was maintained at 26°C. The murine macrophage cell line, J774A.1 (obtained from National Centre for Cell Sciences, Pune) were grown in Dulbecco’s modified Eagle’s medium (DMEM, Gibco, Thermo Fisher Scientific, Waltham, Massachusetts, United States) pH 7.4 supplemented with 2 mM L-glutamine, 100 U/ml penicillin, 100 μg/ml streptomycin, and 10% heat-inactivated FBS at 37°C in a humidified atmosphere containing 5% CO2. Cell number was quantified using a hemocytometer [51].

### Amplification, cloning and sequencing of *LmATP7*

*LmATP7* was PCR-amplified using gene-specific primer P15/16 from intron less *L. major* genomic DNA [52] and cDNA and its expression was checked by running the PCR products on 1% agarose gel. Three primer sets P15/16, P15/17 and P18/19 were used for cloning *LmATP7* into pXG-GFP+ (BamHI and EcoRV sites), pcDNA3.1+ (BamHI and EcoRI sites) and yeast expression vector, p416TEF (as a GFP fusion construct or not) (XbaI and BamHI sites). All four constructs were confirmed by sequencing.

### Transfection

Transfection of DNA into *L. major* was performed using electroporation as described previously [53]. Briefly, 10–30 μg of DNA construct was resuspended in electroporation buffer (21 mM HEPES, 0.7 mM NaH2PO4, 137 mM NaCl and 6 mM glucose; pH 7.4) along with 3.6×10^7^ *L. major* promastigotes. The suspension was incubated in a 0.2 cm electroporation cuvette for 10 min on ice, following which electroporation was performed on the Bio-Rad Gene Pulsar at 450 V, 550 μF capacitance. Transfected cells were selected in appropriate antibiotic-containing medium.

### Generation of *L. major* strain expressing GFP-tagged *LmATP7*

*LmATP7-GFP* construct or vector control (only pXG-GFP) was transfected into wild type *L. major* promastigotes by electroporation as described previously in the transfection section to generate the overexpressing *LmATP7* strain or only GFP expressing strain [54].

### Targeted replacement of *LmATP7* allele of *L. major* with antibiotic-selectable markers

To replace an allele of LmATP7 with NEO gene cassette, we generated the deletion construct in pXG-NEO vector, as described previously [53]. Briefly, 741bp from the 5′ flanking region (starting 174 bp upstream of the start codon) and 740bp from the 3′ flanking region of the *LmATP7* open reading frame were PCR-amplified from *L. major* genomic DNA using primer sets P20/P21 and P22/P23 respectively. HindIII and SalI digested 5′ flanking region, and SmaI and EcoRI digested 3′ flanking region of LmATP7 were ligated on the NEO-encoding open reading frame in pXGNEO plasmid, respectively. The construct was verified by sequencing, following which they were digested with HindIII and BamHI to generate the linearized targeting cassette: 5′LmATP7-NEO-LmATP73′. To generate LmATP7+/− strain, wild-type *L. major* promastigotes were transfected with 10 μg of 5′ LmATP7-NEO-LmATP73′ linearized cassette. LmATP7+/− heterozygous strain was selected and maintained in the presence of 100 μg/ml G418 sulphate.

### Effect of external copper on promastigote growth kinetics

To assess the effect of copper on the growth of *Leishmania* parasites, promastigotes were cultured in M199 medium as mentioned earlier in presence of cupric chloride (CuCl_2_) at a final concentration of 0, 50, 100, 200, 300 or 500 µM. Four different strains of *L. major* including wild type, vector control, LmATP7-OE and LmATP7-Het were used during this study. Parasites were quantified using hemocytometer after 24 hrs, 48 hrs and 72 hrs post incubation in presence of copper to evaluate the growth kinetics. Similarly, to check if copper exerts any toxic effect over macrophage growth, J774A.1 macrophage cultured in DMEM medium was supplemented with CuCl_2_ at a final concentration of 0, 50, 100, 200, 300 or 500 µM. Cell number was measured using hemocytometer following trypan blue staining at 24 hrs and 48 hrs post incubation. Each of the above mentioned studies were performed in triplicate to reach a meaningful statistical analysis.

### Infection of macrophages with *L. major* and estimation of intracellular parasite burden

Infection of J774A.1 murine macrophages with late log-phase *L. major* promastigotes of either wild type, vector control, LmATP7-OE or LmATP7-Het was performed as described previously at a parasite to macrophage ratio of 30:1 [51]. Briefly, J774A.1 macrophage cells were incubated with *L. major* promastigotes for 12 hrs following which the non-phagocytosed parasites were removed and the infection was allowed to continue for 12 hrs, 24 hrs, 30 hrs or 48 hrs. After 12 hrs and 30 hrs time points post infection, cells were harvested to carry out qRT-PCR. After 24 hrs and 48 hrs time points post infection, cells were washed and fixed with acetone-methanol (1:1). Anti-fade mounting medium containing DAPI (VectaShield from Vector Laboratories, Burlingame, CA, United States) was used to stain the nuclei of the fixed infected macrophages. Intracellular parasite burden represented as amastigotes/macrophage cell was quantified by counting the total number of DAPI-stained nuclei of macrophages and *L. major* amastigotes in a field (at least 100 macrophages were counted from triplicate experiments). For measuring the amastigote/ macrophage counts under different concentration of copper treatment, a similar experiment was performed where copper was added post 12 hrs of initial incubation of macrophages with the parasites.

### qRT- PCR of Macrophage and *Leishmania* strains specific copper regulators

Total RNA was isolated from uninfected, wild-type *L. major* infected macrophages as well as from wild-type, *LmATP7* overexpressing and *LmATP7* heterozygous deletion *L. major* strains using TRIzol reagent (Invitrogen, Carlsbad, California, United States). DNA contamination was further removed with DNaseI (Invitrogen, Carlsbad, California, United States) treatment. Verso cDNA synthesis kit (Thermo Fisher Scientific, Waltham, Massachusetts, United States) was used for cDNA preparation from 1 μg of total RNA. Following primers were used for quantification of transcript level of different genes: P1/P2 (*ATP7A*), P3/P4 (*CTR1*), P5/P6 (*CTR2*), P7/P8 (*ATOX1*), P9/P10 (*LmATP7*). Real-time PCR was performed with SYBR green fluorophore (BioRad, Hercules, California, United States) using 7,500 real-time PCR system of Applied Biosystems. The relative transcript level of macrophage specific genes was normalized using wild-type cells as the reference sample and the *β-actin* gene as an endogenous control. In case of LmATP7 mRNA expression, *rRNA45* gene was taken as endogenous control. Amplification of *β-actin* from macrophage and *rRNA45* from *Leishmania* cells was performed using the primer sets P11/P12 and P13/14 respectively. The experiments were performed as per MIQC guidelines.

### Immunofluorescence studies and image analysis

Macrophages were seeded at a density of ∼2×10^5^ cells on glass coverslips and following different experimental steps upto required time points, cells were fixed using acetone: methanol (1:1) for 10 min. Cells were then washed with 1X PBS followed by permeabilization using 0.1% triton-X 100. Both fixation and permeabilization was carried out by keeping it on ice. Cells were then again washed with ice cold 1X PBS, and blocked with 0.2% gelatin for 5 min at room temperature. Incubation with primary antibodies (anti- ATP7A 1:200; anti-Rab11 1:200; anti-Rab7 1:200; anti-Lamp1 1:50 and anti-Lamp2 1:50) was performed for 2 hours at room temperature followed by 1X PBS wash. Cells were reincubated for 1.5 hours at room temperature with either of the following secondary antibodies, goat anti-rabbit Alexa fluor 488 (1:1000) for ATP7A, donkey anti-goat Alexa Fluor 564 (1:1000) for Rab11, goat anti-mouse Alexa fluor 568 (1:1000) for Rab7, Lamp1, Lamp2. Following two 1X PBS washes, coverslips were mounted on glass slides using Fluoroshield™ with DAPI mountant. (#F6057, Sigma-Aldrich, St. Louis, Missouri, United States). All images were visualised with Leica SP8 confocal platform using oil immersion 63X objective and were deconvoluted using Leica Lightning software.

To determine LmATP7 localization in *L. major*, wild type or GFP-only or LmATP7-GFP expressing cells were mounted on poly L-lysine coated coverslips, fixed with acetone: methanol (1:1), and permeabilized with 0.1% Triton X-100 at 4°C. 0.2% gelatine was used to block non-specific binding. Cells were then incubated with either anti-GFP primary antibody or anti- GP63 antibody (1:200) for 1.5 hrs. Thereafter cells were washed with ice-cold PBS and incubated with either goat anti-rabbit Alexa Fluor 488 secondary antibody (1:800) for GFP or with goat anti-mouse Alexa Fluor 568 secondary antibody (1:600) for GP63 for 1.5 hrs in the dark. Post-incubation, cells were washed with PBS and embedded in anti-fade mounting medium containing DAPI. For FM4-64FX staining, LmATP7-GFP expressing cells (under basal or 2hrs of 50µM copper treatment) were incubated for 1min or 10min with the dye prior to the fixation step. Immunostaining was performed using the above-mentioned protocol with primary antibody, anti-GFP (1:200) and secondary antibody, goat anti-rabbit Alexa Fluor Plus 488 (1:800). All images were visualised with Leica SP8 confocal platform using oil immersion 63X objective and were deconvoluted using Leica Lightning software.

### Bioinformatic analysis of LmATP7

The gene (*LmATP7*) for putative copper-transporting ATPase-like protein was identified from TriTrypDB [55]. The SMART server was used to predict transmembrane domains and Pfam [56]. Clustal Omega was used to generate multiple sequence alignment [57]. Multiple sequence alignment visualisation, analysis and editing was performed using Jalview program [58]. ProtParam was used to determine theoretical molecular weight and isoelectric point [59].

### Homology modelling and structural validation

We modelled the three Heavy Metal Binding Domains (HM) of *Leishmania* P-type ATPase copper transporter. The sequence of the three domains that are used for homology modelling include as follows:

HM1: TLNVFGTTCRGCAQHVQENLMTLEGVHSVSVDLDAQLAEVDVDATDAATEFRIEKKMVSMGY
HM2: LLIEGMSCTSCAARIEAKLKQLKGVLGASVNFSAMSGQVLHNPALAPLPKVVSCVADMS
HM3: DGEQPQEGECKCPTNLQAVPVHVGSVMSGYEHRLVVLGMSCASCAARIEHRLRQMPTVLNCTVSFVTGTAV

All these three contain the putative copper binding motif (CXXC), which is thought to be responsible for copper transport. In order to provide a possible structure we performed homology modelling followed by Molecular Dynamics simulations. There is no experimentally resolved 3D structure for the metal binding domains of *Leishmania* ATP7. In order to find the most similar sequence whose structure is known (template) to each of the target sequence, sequence similarity search was performed on the BLASTp online server (http://blast.ncbi.nlm.nih.gov) which were subsequently used as templates for homology modelling performed using the Modeller 9.25 program [60]. This yielded the initial structural guesses for the HMs. The best model was selected with regard to the best Discrete Optimization Protein Energy (DOPE) score [61]. Initially each of the aforementioned best models was solvated by ∼40000 TIP3P water molecules in a box of dimension 80×80×80 Å3 [62]. The physiological concentration (150 mM) of Na+ and Cl- ions along with extra ions were used to neutralize the system. These were subsequently energy minimized using the steepest descent method for 10000 steps, followed by heating it to 300K in 200 ps using Berendsen thermostat and barostat with coupling constant of 0.6 ps. Restraints of 25 kcal/mol/Å2 were applied on heavy atoms during the heating process. Thereafter, equilibration was carried out for 2 ns at constant temperature (300 K) and pressure (1 bar) without any restraints using the same thermostat and barostat with coupling constants of 0.2 ps each. The last 100 ps of NPT simulation was used to calculate the average volume the same, which was used in the final 1000 ns unrestrained NPT equilibration using the Velocity-Rescale thermostat [63] and a Parrinello-Rahman barostat [64] with coupling constant of 0.2 ps. During the simulation, LINCS algorithm was used to constrain all the bonds and Particle Mesh Ewald (PME) method [65] was used for electrostatics. The distance cut-offs for the van der Waals (vdW) and electrostatic long-range interaction was kept at 10 Å. The time step for each simulation was taken to be 2 fs. All of the simulations were performed using molecular dynamics software GROMACS 2019.4 [66] with parameters from the AMBER99SB force field [67].

### Determination of cellular copper concentration by ICP-OES

Wild type and LmATP7-GFP overexpressing *L. major* cells were pelleted down and washed with ice-cold 1X DPBS (Gibco #14200075). Repeated washing was performed (5 times) to ensure there is no trace of copper outside the cells. The pellets were dissolved in DPBS and counted by hemocytometer using formol saline solution. 1×10^6^ cells were digested overnight with 120 µL of 65 % suprapur HNO_3_ at 95^°^C. After digestion, samples were diluted in 6 mL of 1X DPBS and were syringe filtered through a 0.22-micron filter. Copper calibration was done by acid digestion of copper foil (procured from Alfa Aeasar) in 10 mL suprapur HNO_3_ for 1 hr (MWD conditions: Power=400 W; Temperature=1000C; Hold time= 1 h). From the obtained solution, solutions of varying copper strengths were prepared and used for calibration. Copper concentrations of the samples and acid digested DPBS (blank) were determined using a Thermo Scientific Inductively coupled plasma optical emission spectroscopy (ICP-OES) iCAP 6500.

### Strains, media, growth conditions for yeast complementation assay

For yeast complementation studies, *Saccharomyces cerevisiae* BY4742 (wild-type) strain (*MATα his3Δ1 leu2Δ0 lys2Δ0 ura3Δ0*) and an *S. cerevisiae* strain carrying a *CCC2* deletion (ccc2Δ) in the BY4742 background purchased from Euroscarf (Oberursel, Germany) were used. YPD (yeast extract, peptone, and dextrose) medium was used for routinely maintaining both wild-type and deletion strains. For complementation assay, synthetic defined (SD) minimal media containing YNB (yeast nitrogen base), ammonium sulfate, and dextrose supplemented with histidine, leucine, lysine, and methionine (80 mg/liter) were used. Yeast transformations were carried out using lithium acetate method [68]. Human *ATP7B* mRNA was cloned in p416TEF vector (as a positive control) and confirmed by sequencing. *LmATP7*, *GFP-LmATP7* and human *ATP7B* constructs along with empty vector p416TEF were used to transform wild-type (Wt) and ccc2Δ strains. Yeast transformants were selected and maintained on SD medium without uracil (SD-Ura) at 30°C. p416TEF vector contains URA3 selection marker allowing growth in absence of uracil. Wild-type strain was transformed with empty vector to allow its growth on SD-Ura.

### In Vivo Functional Complementation assay in *S. cerevisiae* by dilution spotting

Yeast transformants were grown overnight at 30°C with shaking at 200 rpm in SD-Ura medium. Primary culture was used to inoculate secondary culture in the same selective medium and allowed to grow at 30°C till OD_600_ reached about 0.6. The cells were centrifuged, washed and diluted in sterile water at OD_600_nm = 0.2. Serial dilutions were then made with sterile water (OD_600_ = 0.2, 0.02, 0.002, 0.0002) and 10 μl of cell suspension from each were spotted on plates containing SD-Ura medium with or without 250 μM iron chelator, Ferrozine (Sisco Research Laboratories, Mumbai, India). Plates were incubated at 30°C for 3 days and photographs were taken.

### Statistical analysis

For statistical analysis and plotting, ggplot2 package was used in R v-3.4.0 [69,70]. Statistical analyses were performed by Student’s t test or by Wilcoxon rank-sum test. The results were represented as mean ± SD from minimum three independent experiments. P values of ≤ .05 were considered statistically significant, and levels of statistical significance were indicated as follows: *P ≤ .05, ** P≤ .01, *** P ≤.001, ****P≤ .0001, n.s : non-significant.

## Acknowledgments

This work was supported by DBT-Wellcome Trust India Alliance Fellowship (IA/I/16/1/502369) and Early Career Research Award (ECR/2015/000220) from SERB, Department of Science and Technology (DST), Government of India and IISER K intramural funding to AG and Department of Biotechnology (DBT) and Department of Science and Technology (DST) grants BT/PR21170/MED/29/1109/2016 and EMR/2017/004506, respectively to RD. RP and SM were supported by Pre-doctoral fellowship from Council of Scientific and Industrial Research and SB by DST INSPIRE, India. SS was supported by KVPY fellowship, India. We thank Moulinath Acharya (NIBMG) and Sudipta Chakraborty (NIBMG) for helping us with Real-time PCR and ICP-OES Facility, CCES, IISER-Kolkata for providing us the facility to perform ICP-OES experiment.

## Author contributions

AG, RD, RP, SB and SS designed the experiments and wrote the manuscript. RP, SB, SS and PD did the experiments and analysed the data. SM helped with the statistical analysis. PD and AKB helped with yeast complementation experiments. SS did the homology modelling. All authors reviewed the results and approved the final version of the manuscript.

**The authors declare no competing financial interests.**

## Supplementary figs

**S1.**
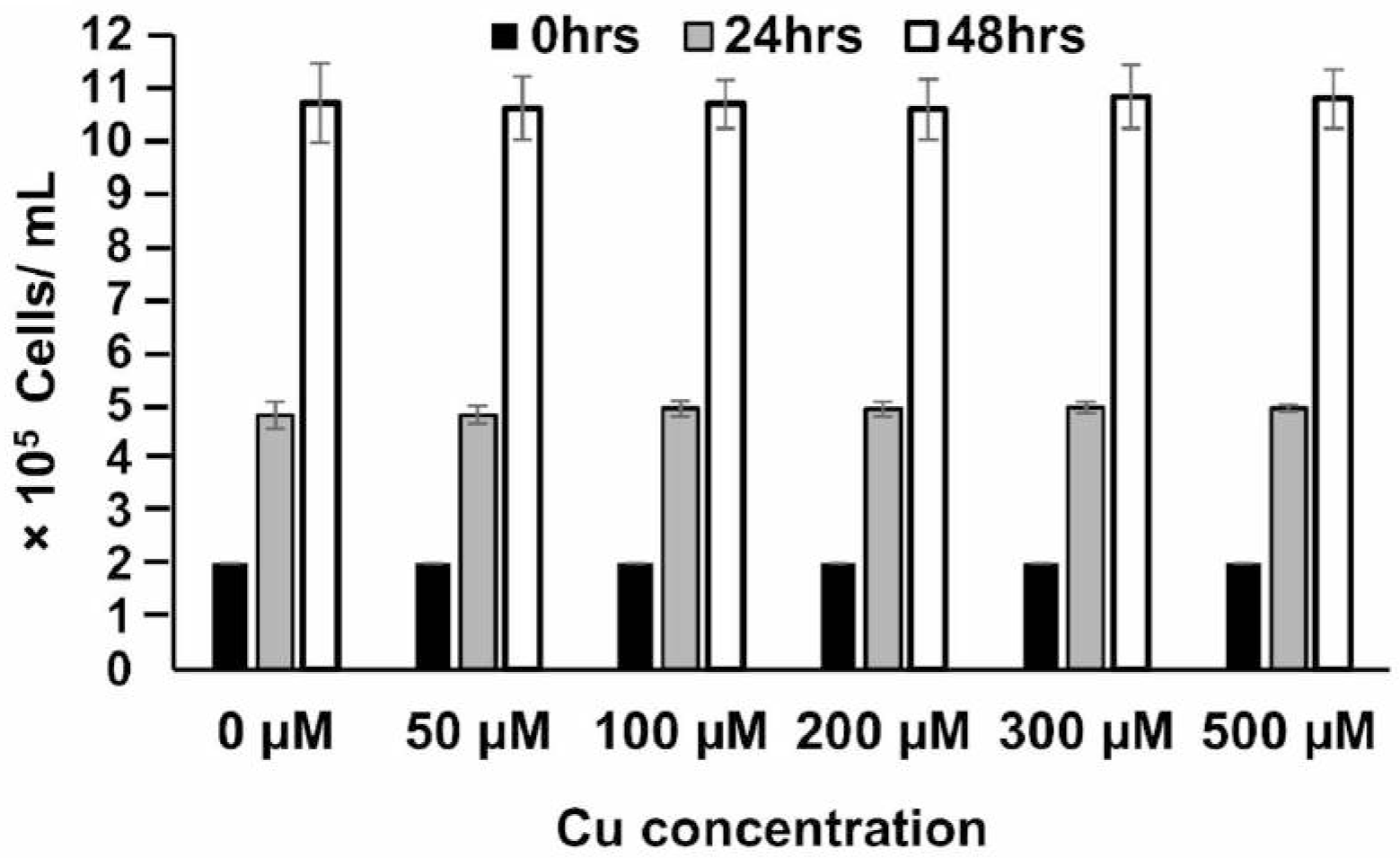
J774A.1 macrophage cell growth in presence of external copper. Growth of J774A.1 macrophage cells were measured in media supplemented with 0- 500µM of copper for 0- 48 hrs. Initially, 2.0×10^6^ cells were added to the media (0 hrs) and further at the above mentioned time-points post incubation, cells were counted using trypan-blue exclusion method by haemocytometer-based cell counting. Error bars represent mean ± SD of values from three independent experiments.

**S2.**
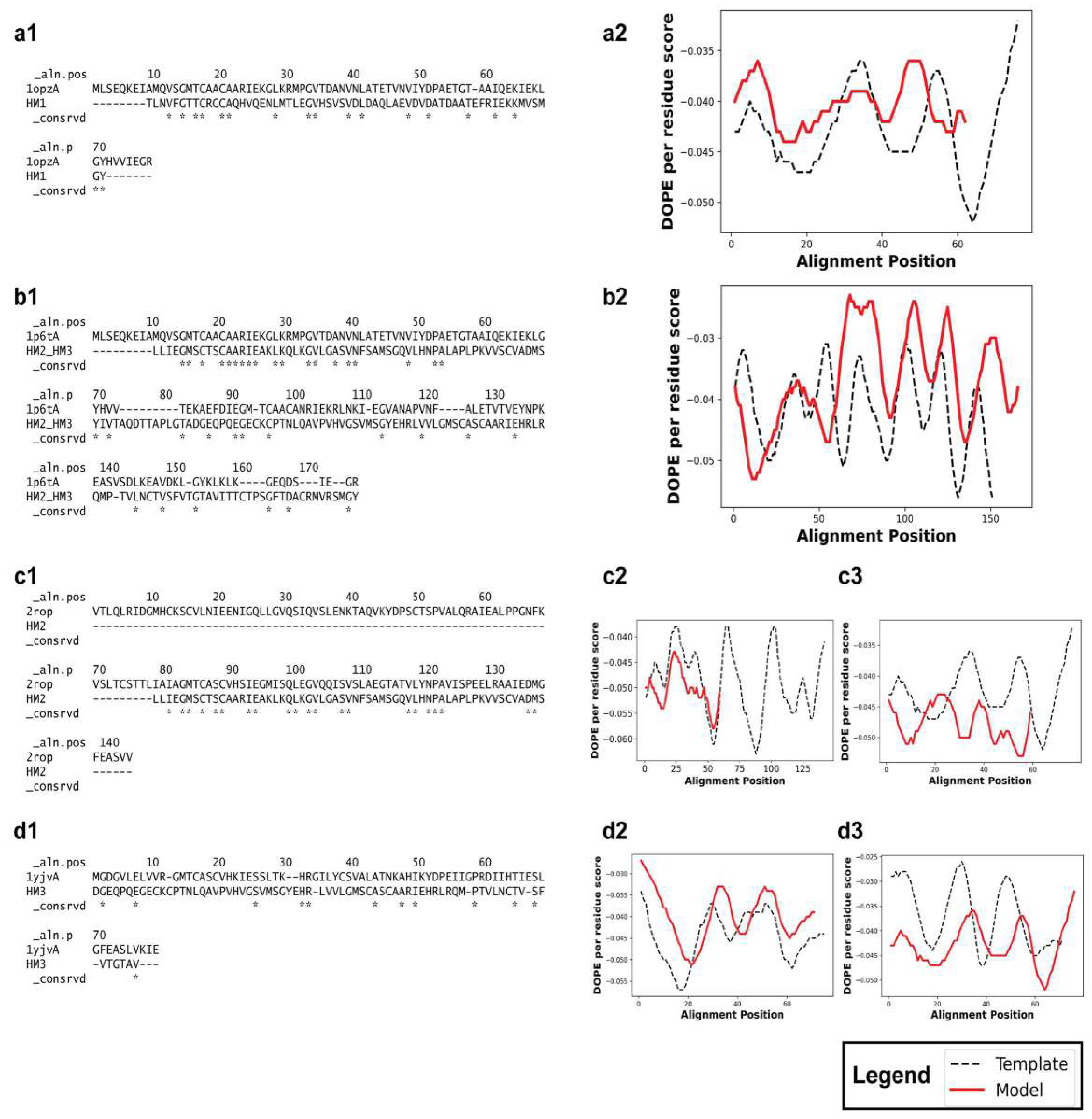
Homology model of the Copper binding motifs of LmATP7 protein. **(a1)** Sequence comparison between HM1 and 1OPZ_A chain with the conserved residues being indicated by “*” symbol. **(a2)** DOPE-per-residue score versus Alignment Position plotted for HM1 (model) and template (1OPZ.pdb). **(b1)** Sequence comparison between combined HM2 and HM3 and 1P6T_A chain with the conserved residues being indicated by “*” symbol. **(b2)** DOPE per residue score versus Alignment Position plotted for HM2_HM3 (model) and template (1P6T.pdb)(c1) Sequence comparison between combined HM2 and 2ROP_A chain with the conserved residues being indicated by “*” symbol. **(c2, c3)** DOPE per residue score versus Alignment Position plotted for HM2 (model) and template **(b)** 2ROP.pdb) and **(c)** 1OPZ.pdb). The third plot shows, with the template corresponding to *B. subtilis* CopA shows clear mismatch between the template and model while the second plot with the template corresponding to metal binding domain 4 of ATP7B shows a greater match. **(d1)** Sequence comparison between combined HM3 and 1YJV_A chain with the conserved residues being indicated by “*” symbol. **(d2,d3)** DOPE per residue score versus Alignment Position plotted for HM3 (model) and template **(b)** 1YJV.pdb) and **(c)** 1OPZ.pdb). The third plot shows, with the template corresponding to *B. subtilis* CopA shows clear mismatch between the template and model while the second plot with the template corresponding to metal binding domain 6 of ATP7A shows a greater match.

**S3.**
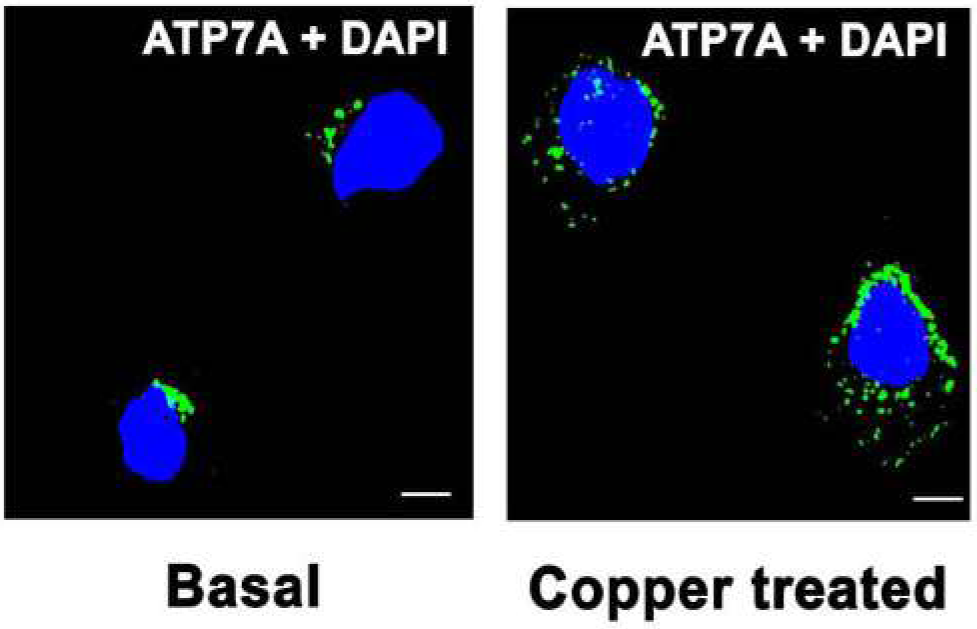
Control (Basal) and copper treated (50µM; 2 hrs) macrophages immunostained with anti-ATP7A (green). Representative confocal microscopy images represent the localization of ATP7A (green) under basal and copper treatment. Nuclei were stained with DAPI (blue). Scale bar: 5µm.

**S4.**
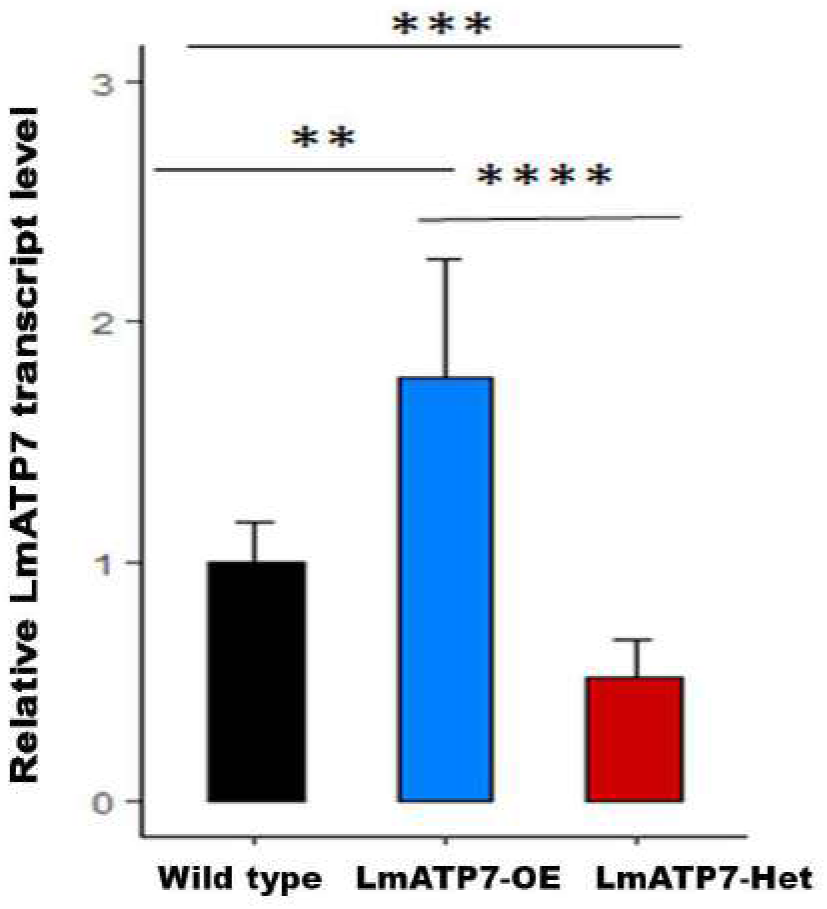
Real time PCR confirms the generation of LmATP7 overexpressed and heterozygous deletion *L. major.* Bar diagram showing *LmATP7* transcript level normalized against *rRNA45* in Wild type (black bar), LmATP7 overexpressed (LmATP7-OE, blue bar) and LmATP7 heterozygous deletion (LmATP7-Het, red bar) *L. major*. Error bars represent mean ± SD of values calculated from three independent experiments. * indicate significant difference with respect to the *LmATP7* transcript level. *P≤0.01, ***P≤0.001, ****P≤0.0001; (Student’s t-test).

**S5.**
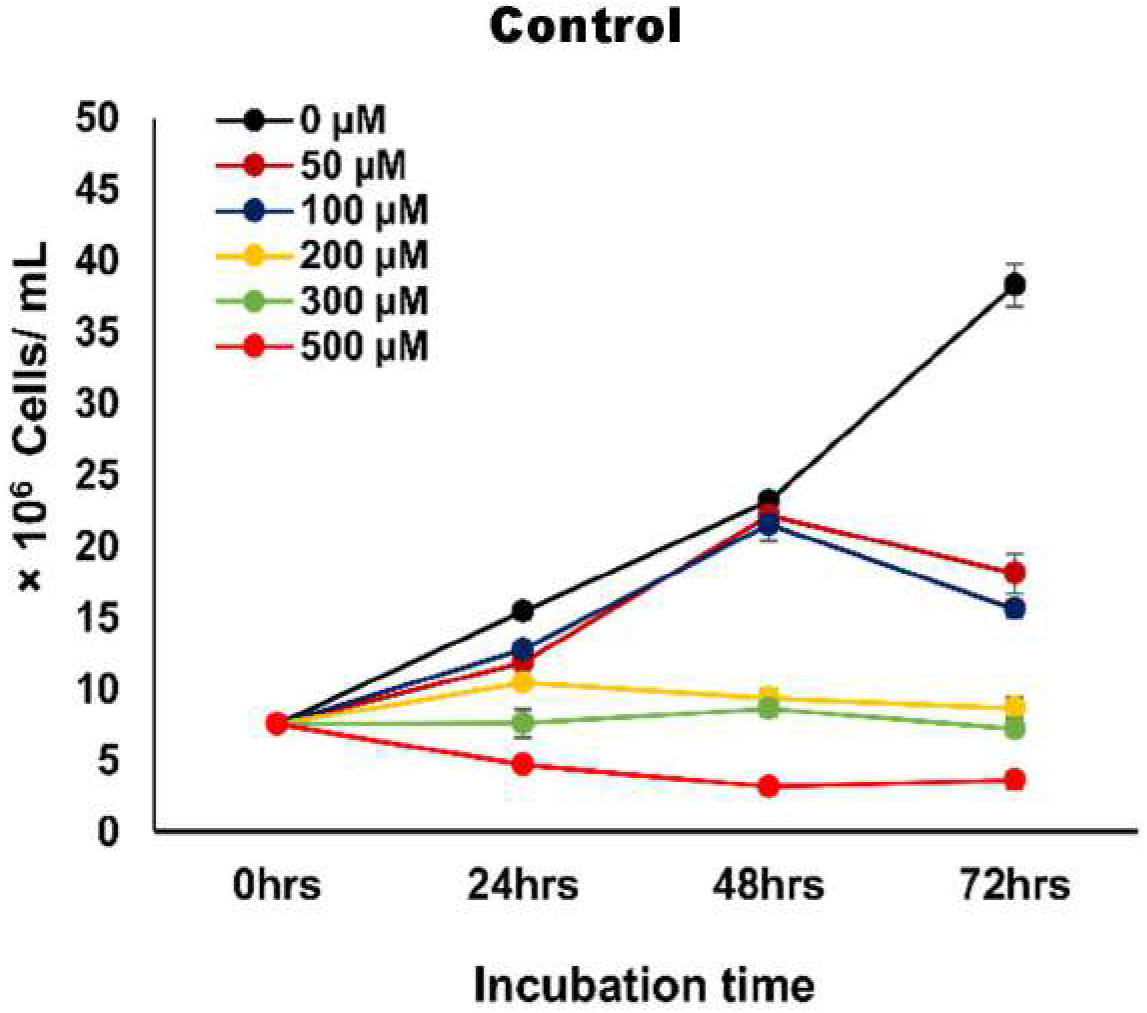
Growth kinetics of vector control *L. major* promastigotes in presence of external copper. pXG-GFP vector control *L. major* cells were grown in 0- 500µM externally added copper concentration for 24 hrs, 48 hrs and 72 hrs. Initially, 7.5×10^6^ cells were added to the media at time point 0 hrs and further at the above mentioned time-points cells were counted by haemocytometer-based cell counting. Error bars represent mean ± SD of values from three independent experiments.

**S6.**
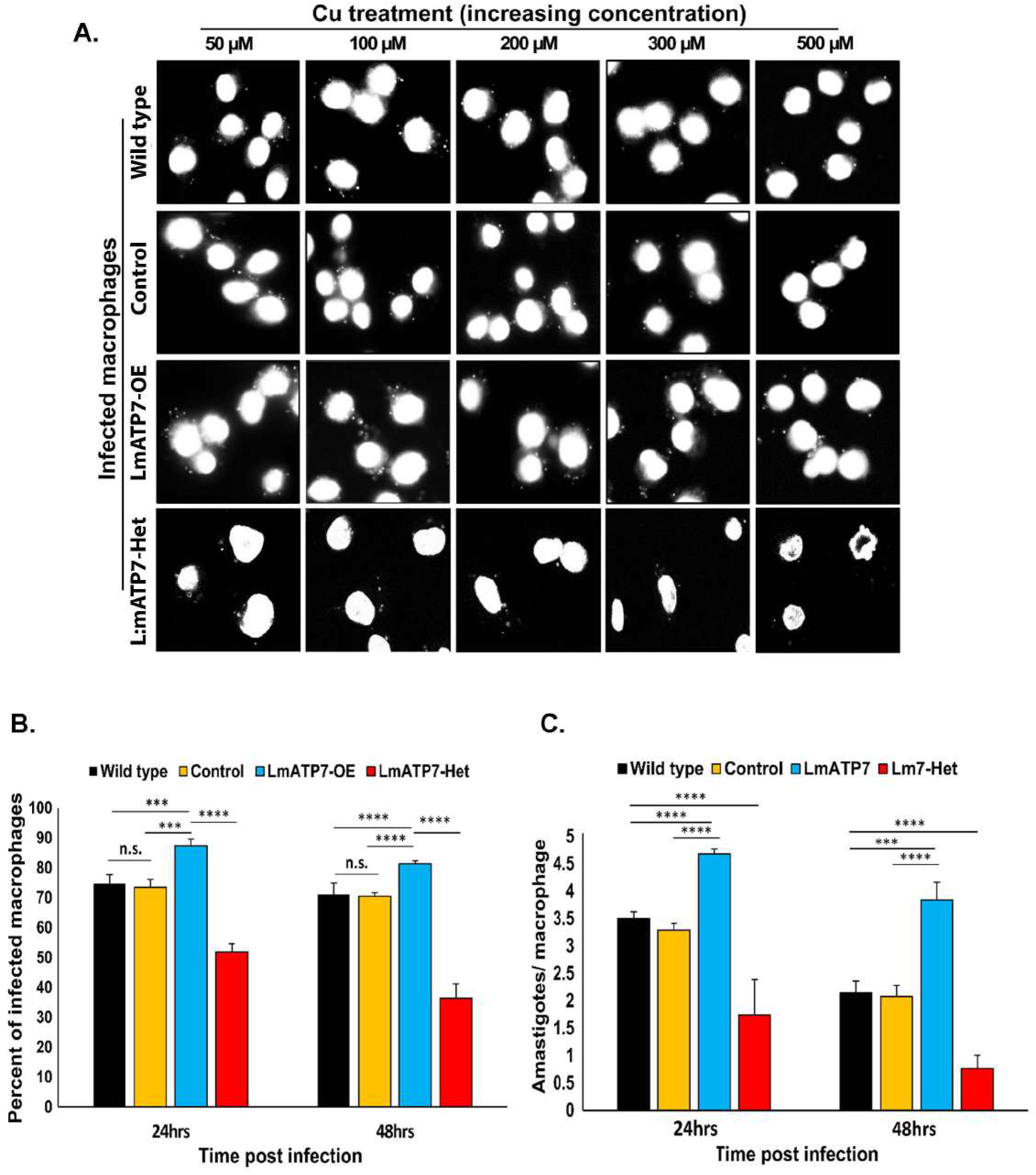
LmATP7 gene is important for the survival and infectivity of *L. major*. **(A)** Representative DAPI stained epifluorescence microscopy images of three independent experiments showing J774A.1 macrophage infected with Wild type, vector control (Control), LmATP7 overexpressed (LmATP7-OE) or LmATP7 heterozygous deletion (LmATP7-Het) *L. major* strains in presence of external copper (50-500µM) at 24 hrs post infection. In the images larger nuclei represent macrophage cells which are surrounded by smaller nuclei of *Leishmania* amastigotes. Images were taken under Olympus IX-81 epifluorescence microscope at 60X magnification. **(B)** J774A.1 macrophage cells were infected with Wild type, Control, LmATP7-OE or with LmATP7-Het *L. major* promastigotes for either 24 or 48 hrs in absence of external Cu. Cells were stained with DAPI and visualized under Olympus IX-81 epifluorescence microscope. The percent of infected macrophages (no. of infected macrophage cells/ 100 macrophages) was quantified by counting the total number of DAPI-stained nuclei of infected macrophages in a field. At least 100 macrophages were counted from triplicate experiments. Error bars represent mean± SD of values calculated from at least three independent experiments. (C) Graphical representation of amastigotes/ macrophage count at 24 hrs and 48 hrs post infection. J774A.1 macrophages were infected with Wild type (black bar), Control (yellow bar), LmATP7-OE (blue bar) or LmATP7-Het (red bar) strains in absence of external Cu. At least 100 macrophages were counted from triplicate experiments. Error bars represent mean ± SD of values from three independent experiments. * indicates the significance of difference in amastigote/ macrophage count in case of Control, LmATP7-OE and LmATP7-Het strains compared to Wild type cells at a particular time point. *P≤0.05, **P≤0.01, ***P≤0.001, ****P≤0.0001; (Student’s t-test).

