## Supplemental Table 1 for "The novel leishmanial Copper P-type ATPase plays a vital role in intracellular parasite survival" (from another version of this preprint: version 2, 2021-01-02)

Table S1. Oligonucleotides used in this paper

| Primer | Primer Sequence (5'- 3') |
|---|---|
| P1 | GAAGAGGTCGGACTGCTGTC |
| P2 | CCTTAGTAATGCCAACCTGAGAAGC |
| P3 | CAAGATAGCCCGAGAGGGTC |
| P4 | GATGTGCAGCACTGTCTGC |
| P5 | AGATGAGGCCGTGCTTCTC |
| P6 | CCTGGTCTGGTCCCAAGATG |
| P7 | GCACGAGTTCTCCGTGGAC |
| P8 | GGGCCAAGGTAGGAAACAGC |
| P9 | TGTCATCCGGTATCTGCACA |
| P10 | CGGCGTACACATCTTTCGAG |
| P11 | GGCTGTATTCCCCTCCATCG |
| P12 | CCAGTTGGTAACAATGCCATG |
| P13 | CCTACCATGCCGTGTCCTTCTA |
| P14 | AACGACCCCTGCAGCAATAC |
| P15 | CGCGGATCCATGGGCGCGACGACT |
| P16 | CGAGATATCTGACATGAAATAGAAATCGCGTT |
| P17 | CCGGAATTCTCACATGAAATAGAAATCGCGTT |
| P18 | CGCGGATCCATGGGCGCGACGACTGC |
| P19 | CCCAAGCTTTCACATGAAATAGAAATC |
